# Combining computational controls with natural text reveals new aspects of meaning composition

**DOI:** 10.1101/2020.09.28.316935

**Authors:** Mariya Toneva, Tom M. Mitchell, Leila Wehbe

## Abstract

To study a core component of human intelligence—our ability to combine the meaning of words—neuroscientists have looked to theories from linguistics. However, linguistic theories are insufficient to account for all brain responses that reflect linguistic composition. In contrast, we adopt a data-driven computational approach to study the combined meaning of words beyond their individual meaning. We term this product “supra-word meaning” and investigate its neural bases by devising a computational representation for it and using it to predict brain recordings from two imaging modalities with complementary spatial and temporal resolutions. Using functional magnetic resonance imaging, we reveal that hubs that are thought to process lexical-level meaning also maintain supra-word meaning, suggesting a common substrate for lexical and combinatorial semantics. Surprisingly, we cannot detect supra-word meaning in magnetoencephalography, which suggests the hypothesis that composed meaning might be maintained through a different neural mechanism than the synchronized firing of pyramidal cells. This sensitivity difference has implications for past neuroimaging results and future wearable neurotechnology.

---

Understanding language in the real-world requires us to compose the meaning of individual words in a way that makes the final composed product more meaningful than the string of isolated words. For example, we understand the statement that “Mary finished the apple” to mean that Mary finished *eating* the apple, even though “eating” is not explicitly specified (Pylkkänen, 2020). This *supra-word meaning*, or the product of meaning composition beyond the meaning of individual words, is at the core of language comprehension, and its neurobiological bases and processing mechanisms must be specified in the pursuit of a complete theory of language processing in the brain.

Different types of supra-word meaning exist. In addition to coercion, such as “finished eating” in “Mary finished the apple” (Pylkkänen, 2020), other examples of supra-word meaning include: a specific contextualized meaning of a word or phrase (e.g. “green banana” evokes the meaning of an unripe, rather than simply green-colored, banana) that can also distinguish between different senses of the same word (e.g. “play a game” versus “theater play”), and the different meaning of two events that can be described with the same words but reversed semantic roles (e.g. “John gives Mary an apple” and “Mary gives John an apple”). Previous works have focused on specific types of supra-word meaning in carefully controlled experiments (Pylkkänen and McElree, 2007; Baggio *et al*., 2010; Bemis and Pylkkänen, 2011; Brooks and Cid de Garcia, 2015; Kim and Pylkkänen, 2019). However, much is left to know about the brain processing of supra-word meaning in naturalistic language. For instance, in order to understand how meaning is processed and composed and how the brain makes sense of language, knowing where the supra-word meaning is maintained is an essential requirement. One proposed hypothesis is that the ventro-medial prefrontal cortex is the possible area that represents the product of meaning composition (Pylkkänen, 2020).

How can we find the regions that represent supra-word meaning? One approach is to focus on every type of supra-word meaning and design a controlled experiment to study it. This approach is not without challenges as it would require a large number of experimental conditions, and carefully balancing the condition of interest and the control condition for all types of supra-word meaning might be challenging. One alternative is to use a complex natural text that readily contains various types of supra-word meaning and build representations that characterize this set of supra-word meanings. Deep neural network language models are the current most powerful tools for building such representations (Peters *et al*., 2018; Devlin *et al*., 2018; Brown *et al*., 2020). Though these NLP systems are not specifically designed to mimic the processing of language in the brain, representations of language extracted from these NLP systems have been shown to predict the brain activity of a person comprehending language better than ever before (Wehbe *et al*., 2014a; Jain and Huth, 2018; Toneva and Wehbe, 2019; Schrimpf *et al*., 2020; Caucheteux and King, 2020; Goldstein *et al*., 2021).

After being trained to predict a word in a specific position from its context on extremely large corpora of text, neural network language models achieve unprecedented performance on various natural language processing (NLP) tasks(Peters *et al*., 2018; Devlin *et al*., 2018; Brown *et al*., 2020). One can use these models to extract representations for the meaning of stimulus text. While these representations are often difficult to interpret (we don’t know what the dimensions of the representational space correspond to, what processes are being computed, or what type of composition is happening at a given time), they can still help us achieve our goal of identifying the regions that represent supra-word meaning. This is because the neural network representations could be assumed to contain at least some of the aspects of supra-word meaning (otherwise they would fail at many of the NLP tasks(Levesque *et al*., 2012; Marvin and Linzen, 2018)). If we can isolate the information in the neural network representations of a sequence of words that is not contained in the individual words themselves, then we would be isolating some aspects of supra-word meaning. We can then identify brain regions that are well predicted by this isolated supra-word meaning.

We thus build a computational object for supra-word meaning using neural network representations, which we call a supra-word embedding. Specifically, we construct the following computational representation of supra-word meaning: a “supra-word embedding” is the part of a contextualized word embedding extracted from the NLP system that is orthogonal to the individual word meanings that make-up the adjacent context. This computational object acts as a set that includes some types of supra-word meaning and that enables us to investigate brain representations and identify regions that encode supra-word meaning. The current work relies on deep learning models because of their expressivity and ability to predict brain activity in naturalistic contexts, which at the moment is not matched by methods that are tied to linguistic theory. Linguistic theory has traditionally focused on a different set of problems than the problem of predicting words and word sequences with high accuracy across a large variety of contexts, and is well equipped to shed light on more systematic features of language (Baroni, 2021). While our analysis relies on deep learning models, it does not exclude future analyses that are more closely tied to linguistic theory. Future work can indeed focus on constructing representations for specific types of supra-word meaning, or on interpreting the contents of supra-word embeddings to identify which types of supra-word meaning they contain.

We study the neural bases of supra-word meaning by using its computational representation “supra-word embedding” and data from naturalistic reading in two neuroimaging modalities. We find that the supra-word embedding predicts functional magnetic resonance imaging (fMRI) activity in the anterior and posterior temporal cortices, suggesting that these areas represent composed meaning. The posterior temporal cortex is considered to be primarily a site for lexical (i.e. word-level) semantics (Hagoort, 2020; Hickok and Poeppel, 2007) so our finding that it also maintains supra-word meaning suggests a common substrate for lexical and combinatorial semantics. Furthermore, we find clusters of voxels in both the posterior and anterior temporal lobe that share a common representation of supra-word meaning, suggesting the two areas may be working together to maintain the supra-word meaning. We replicate these findings in an independent fMRI dataset recorded from a different experimental paradigm and a different set of participants.

The second neuroimaging modality that we utilize to investigate the neural bases of supra-word meaning is magnetoencephalography (MEG). While MEG and fMRI are both thought to be primarily driven by post-synaptic cellular processes and many traditional localization studies have found similar location of activations in fMRI and MEG for the same task, the relationship between these two modalities is complex and still not fully understood (Hall *et al*., 2014). For example, some discordances have been observed in the primary cortex, where MEG has sensitivity to spatial frequency (Muthukumaraswamy and Singh, 2008, 2009) and color (Swettenham *et al*., 2013) while fMRI does not (Muthukumaraswamy and Singh, 2008, 2009; Swettenham *et al*., 2013). At the same time, fMRI-MEG fusion, a method that relates the representations of the same stimuli in both modalities, has identified regions that are sensitive to the properties of written individual words when measured in fMRI but without a significant correspondence in MEG (Leonardelli and Fairhall, 2022). We also find a mismatch between our MEG and fMRI results. We find that it is very difficult to detect the representation of supra-word meaning in MEG activity. MEG has been shown to reveal signatures of the *computations* involved in incorporating a word into a sentence (Halgren *et al*., 2002; Lyu *et al*., 2019), which are themselves a function of the composed meaning of the words seen so far. However, our results suggest that the sustained *representation* of the composed meaning may rely on neural mechanisms that do not lead to reliable MEG activity. This hypothesis calls for a more nuanced understanding of the body of literature on meaning composition and has important implications for the future of brain-computer interfaces.

## Results

### Computational controls of natural text

We built on recent progress in NLP that has resulted in algorithms that can capture the meaning of words in a particular context. One such algorithm is ELMo (Peters *et al*., 2018), a powerful language model with a bi-directional Long Short-Term Memory (LSTM) architecture. ELMo estimates a *contextualized* embedding for a word by combining a *non-contextualized* fixed input vector for that word with the internal state of a forward LSTM (containing information from previous words) and a backward LSTM (containing information from future words). To capture information about word *t*, we used the input vector for word *t*. To capture information about the context preceding word *t*, we used the internal state of the forward LSTM computed at word *t* – 1 (Fig. 1B). We did not include information from the backward LSTM, since it contains future words which have not yet been seen at time *t*. Note that ELMo’s forward and backward LSTMs are trained independently and do not influence one another (Peters *et al*., 2018). We have also experimented with GPT-2 (Radford *et al*., 2019), which is a more complex language model with a transformer-based architecture and 12 internal layers, and observed that our findings from ELMo replicate (see Results). We choose to focus on ELMo because of its good performance at language tasks and at predicting brain recordings (Toneva and Wehbe, 2019), and yet relative simplicity with respect to other recent language models.

**Figure 1:**
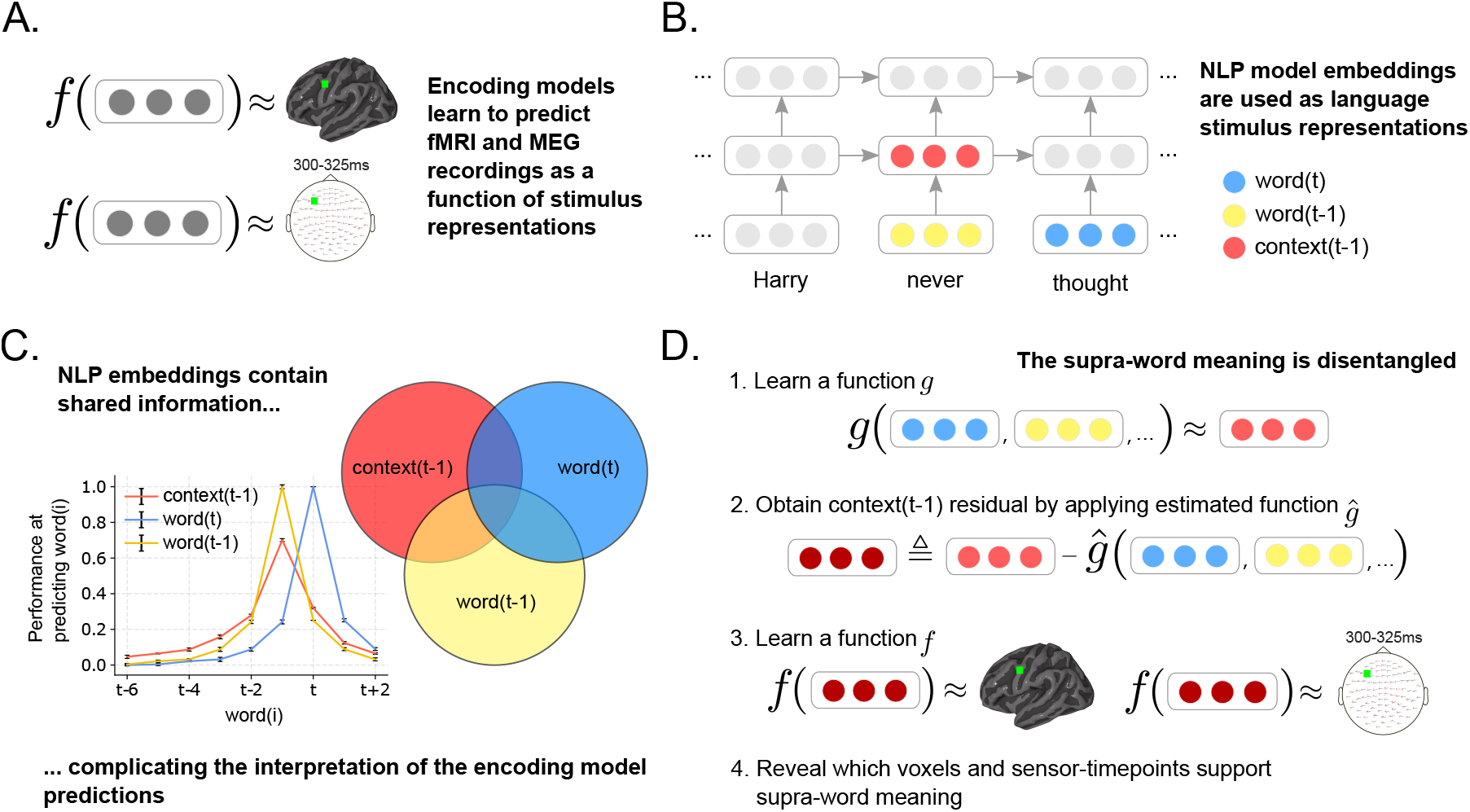
Approach. **(A)** An encoding model *f* learns to predict a brain recording as a function of representations of the text read by participant during the experiment. A different function is learned for each voxel in fMRI and sensor-timepoint in MEG. **(B)** Stimulus representations are obtained from an NLP model that has captured language statistics from millions of documents. This model represents words using context-free embeddings (shown in yellow and blue) and context embeddings (shown in red). Context embeddings are obtained by continuously integrating each new word’s context-free embedding with the most recent context embedding. **(C)** Context and word embeddings share information. The performance of the context and word embeddings at predicting the words at surrounding positions is plotted for different positions. The context embedding contains information about up to 6 past words, and word embeddings contains information about embeddings of surrounding words. To isolate the representation of supra-word meaning, it is necessary to account for this shared information. **(D)** Supra-word meaning is modeled by obtaining the residual information in the context embeddings after removing information related to the word embeddings. We refer to this residual as the *supra-word embedding* or *residual context embedding*. The supra-word embedding is used as an input to an encoding model f, revealing which fMRI voxels and MEG sensor-timepoints are modulated by supra-word meaning.

To study supra-word meaning, the meaning that results from the composition of words should be isolated from the individual word meaning. ELMo’s context embeddings contain information about individual words (e.g., ’finished’, ’the’, and ’apple’ in the context ’finished the apple’) in addition to the implied supra-word meaning (e.g., eating) (Fig. 1C). In fact, ELMo’s context embedding of a word t is strongly linearly related to the next word t+1, current word t, and the previous word t-1 (see Suppl. Fig. S1). We post-processed the context embeddings produced by ELMo to remove the contribution due to the context-independent meanings of individual words. We constructed a “residual context embedding” by removing the shared information between the context embedding and the meanings of the individual words (Fig. 1D, also Suppl. Fig. S1).

To investigate the neural substrates and temporal dynamics of supra-word meaning, we trained encoding models, as a function of the constructed residual context embedding, to predict the brain recordings of nine fMRI participants and eight MEG participants as they read a chapter of a popular book in rapid serial visual presentation. We further replicate our fMRI findings in a second fMRI dataset with a different experimental paradigm that was collected while 6 participants viewed a popular movie. The encoding models predict each fMRI voxel and MEG sensor-timepoint, from the text read by the participant up to that time point (Fig. 1A). The prediction performance of these models was tested by computing the correlation between the model predictions and the true held-out brain recordings. Hypothesis tests were used to identify fMRI voxels and MEG sensor-timepoints that were significantly predicted by the residual context embedding. For more details about the training procedure and hypothesis tests, see Materials and Methods.

### Detecting regions that are predicted by supra-word meaning

To identify brain areas that represent supra-word meaning, we focus on the fMRI portion of the experiment. We find that many areas previously implicated in language-specific processing (Fedorenko *et al*., 2010; Fedorenko and Thompson-Schill, 2014) and word semantics (Binder *et al*., 2009) are significantly predicted by the full context embeddings across subjects (voxel-level permutation test, Benjamini-Hochberg FDR control at 0.01 (Benjamini and Hochberg, 1995)). These areas include the bilateral posterior and anterior temporal cortices, angular gyri, inferior frontal gyri, posterior cingulate, and dorsomedial prefrontal cortex (Fig. 2A and Suppl. Fig. S2 and S3). A subset of these areas is also significantly predicted by residual context embeddings. To quantify these observations, we select regions of interest (ROIs) based on the works above (Fedorenko *et al*., 2010; Binder *et al*., 2009), using ROI masks that are entirely independent of our analyses and data (see Materials and Methods). Full context embeddings predict a significant proportion of the voxels within each ROI across all 9 participants (Fig. 2B; ROI-level Wilcoxon signed-rank test, *p* < 0.05, Holm-Bonferroni correction (Holm, 1979)). In contrast, residual context embeddings predict a significant proportion of only the anterior and posterior temporal lobes. While the full context embedding is predictive of much of the fMRI recordings in the language and semantic networks, the the residual context embedding is more selectively predictive of two language regions – the anterior (ATL) and posterior temporal lobes (PTL). These results are not specific to word representations obtained from ELMo. Using a different language model–GPT-2–to obtain the full and residual context embeddings replicates the results using ELMo (see Materials and Methods), showing that all bilateral language ROI are predicted significantly by the full context embedding, and that the bilateral ATL and PTL are predicted significantly by the residual context embedding (see Suppl. Fig. S6).

**Figure 2:**
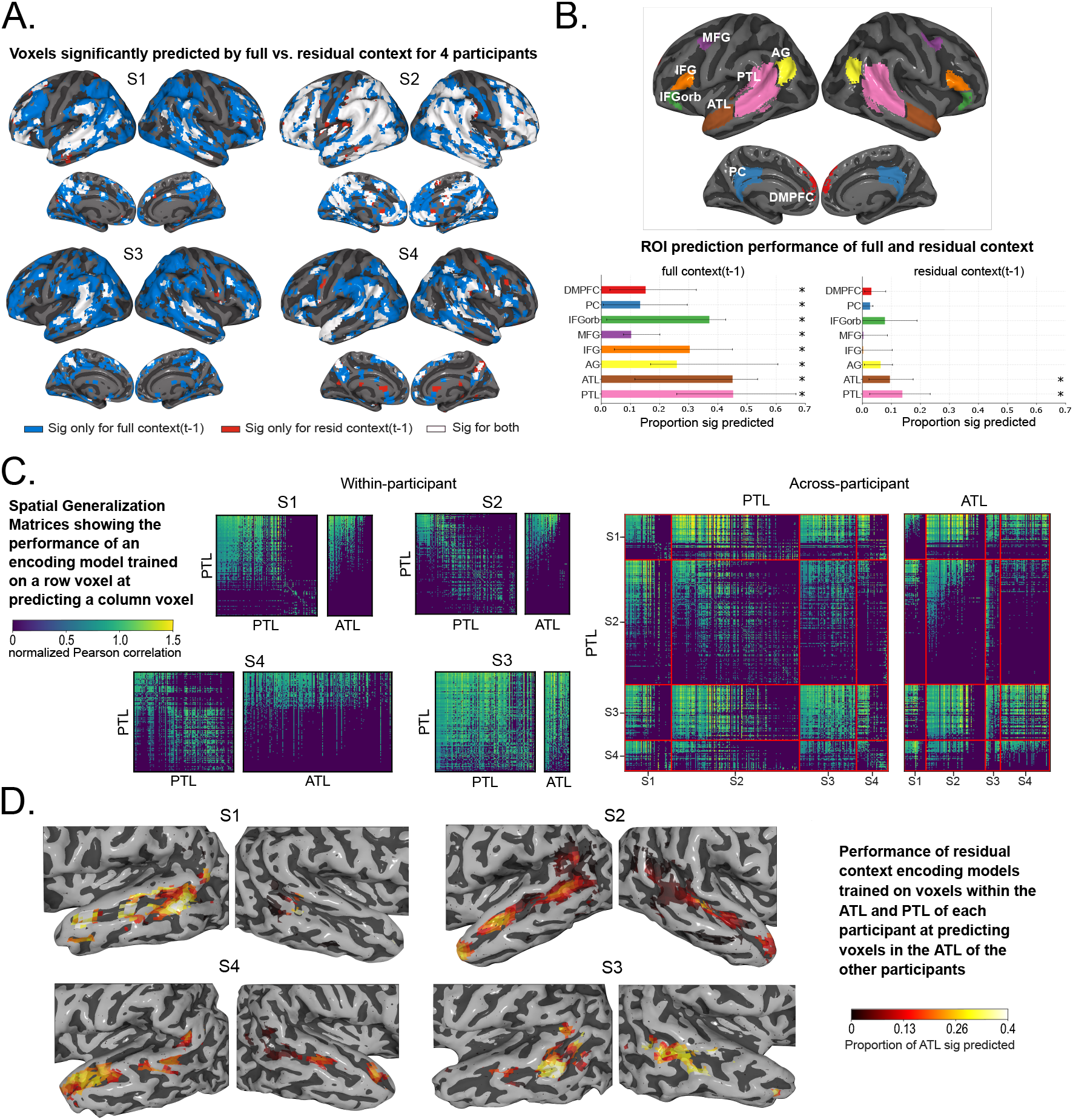
fMRI results. Visualizations for 4 of 9 participants with remainder available in Suppl. Fig. S3-S5. Voxel-level significance is FDR corrected at *α* = 0.01. **(A)** Voxels significantly predicted by full-context embeddings (blue), residual-context embeddings (red), or both (white), visualized in MNI space. Most of the temporal cortex and IFG is predicted by full context embeddings, with residual context embeddings mostly predicting a subset of those areas. **(B)** ROI-level results. (Top) Language system ROIs (Fedorenko *et al*., 2010) and two semantic ROIs (Binder *et al*., 2009). (Bottom) Proportion of ROI voxels significantly predicted by (Left) full context and (Right) residual context embeddings. Displayed are the median proportions across all participants and the medians’ 95% confidence intervals. Full context predicts all ROIs (ROI-level Holm-Bonferroni correction, *p* < 0.05), while residual context predicts only bilateral ATL and PTL. **(C)** Spatial Generalization Matrices. Models trained to predict PTL voxels are used to predict PTL and ATL voxels (within-participant (Left), and across-participants (Right)). PTL cross-voxel correlations form two clusters: models that predict activity for voxels in one cluster can also predict activities of other voxels in the same cluster, but not activities for voxels in the other cluster. Across participants, only one of these clusters has voxels that predict ATL voxels. **(D)** Performance of models trained on ATL and PTL voxels at predicting other participants’ ATL. All participants show a cluster of predictive voxels in the pSTS.

Do the parts of the ATL and PTL that are predicted by supra-word meaning process the same information? Inspired by temporal generalization matrices (King and Dehaene, 2014), we introduce *spatial generalization matrices* that estimate the pairwise similarity of voxel representations (see Materials and Methods). The spatial generalization matrices reveal that the PTL can be divided into two main clusters such that the models of voxels in one cluster can also predict other voxels in that cluster but not in the other cluster (Fig. 2C and Suppl. Fig. S4; voxel-level permutation test, Benjamini-Hochberg FDR controlled at level 0.01). Furthermore, the models of voxels within one of the PTL clusters, but not the other, significantly predict voxels in the ATL. The division of the PTL into two clusters, one of which is predictive of the ATL, can be observed within- (Fig. 2C, left), and across-participants (Fig. 2C, right). In contrast, the ATL voxels show only one cluster of voxels that are predictive both of other ATL voxels and also of PTL voxels (Suppl. Fig. S4). This pattern indicates that the organization of information in the ATL and parts of the PTL is shared and consistent across participants. To localize this shared representation, we visualize how well each ATL and PTL voxel predicts the other participants’ ATLs (Fig. 2D and Suppl. Fig. S5). ATL voxels are predictive of significant proportions of the ATL across participants, reinforcing the single cluster of ATL voxels observed in the spatial generalization matrices. Much of the left PTL predicts a significant proportion of the ATL across participants, whereas much of the right PTL does not (ROI-level Wilcoxon signed-rank test, *p* < 0.05, Holm-Bonferroni correction). The left PTL appears further subdivided, with a cluster of voxels in the posterior Superior Temporal Sulcus (pSTS) being more predictive. This suggests that the ATL and the left pSTS process a similar facet of supra-word meaning.

To test whether the findings that the ATL and PTL support supra-word meaning are specific to our dataset or experimental paradigm, we conducted a replication analysis using a second fMRI dataset. In this second fMRI dataset, acquired by the Courtois NeuroMod Group, participants viewed a full-length popular movie and the computational representation of supra-word meaning (i.e. the residual context embedding) was computed based on the speech in the movie. We find that the results from the naturalistic reading paradigm that the residual context embedding predicts most significantly the bilateral ATL and PTL are repeated with this dataset (see Suppl. Fig. S7 for ROI-level results, Suppl. Fig. S8 for group-level voxel-wise significance masks, and Suppl. Fig. S9 for individual voxel-wise significance masks). In addition to the bilateral ATL and PTL, the residual context embedding significantly predicts the inferior frontal gyrus (IFG) orbitalis and the posterior cingulate at a FDR-corrected pvalue of 0.045. We further also see that the ATL and a portion of the left PTL are predicted by a similar facet of supra-word meaning (Suppl. Fig. S11,S4). These results replicate our findings about the bilateral ATL and PTL being significantly predicted by the supra-word meaning computational representation, which is encouraging because the two fMRI datasets were collected under completely different paradigms, different sensory modalities (reading vs. listening), and different participant population.

### The processing of supra-word meaning is invisible in MEG

To study the temporal dynamics of the emergence and representation of supra-word meaning, we turn to the MEG portion of the experiment (Fig. 3). We computed the proportion of sensors that are significantly predicted at different spatial granularity – the whole brain (Fig. 3A), by lobe subdivisions (Suppl. Fig. S12), and finally at each sensor neighborhood location (Fig. 3B; sensor-timepoint level permutation test, Benjamini-Hochberg FDR control at *α* = 0.01, see Suppl. Fig. S14 for sensor-level results for individual participants). The full context embedding is significantly predictive of the recordings across all lobes (Fig. 3A, performance visualized in lighter colors; timepoint-level Wilcoxon signed-rank test, *p* < 0.05, Benjamini-Hochberg FDR correction). Surprisingly, we find that the residual context does not significantly predict any timepoint in the MEG recordings at any spatial granularity. This surprising finding leads to two conclusions. First, supra-word meaning is invisible in MEG. Second, what is instead salient in MEG recordings is information that is shared between the context and the individual words. These results are not specific to word representations obtained from ELMo. Using a different language model–GPT-2–to obtain the full and residual context embeddings replicates the results using ELMo (see Materials and Methods), showing that while the full context embedding predicts the MEG recordings significantly, the residual context embedding fails to significantly predict any sensor-timepoint across participants (see Suppl. Fig. S15). Further, these results repeated in data from one subject listening to 70 minutes of spoken stories (totalling 15030 words) from Huth *et al*. (2016) (Huth *et al*., 2016), shown in Suppl. Fig. S16. We followed a similar analysis with this data to reveal that while the full context embedding is well predictive of many sensor-timepoint combinations, the residual context is not predictive at any sensor or timepoint.

**Figure 3:**
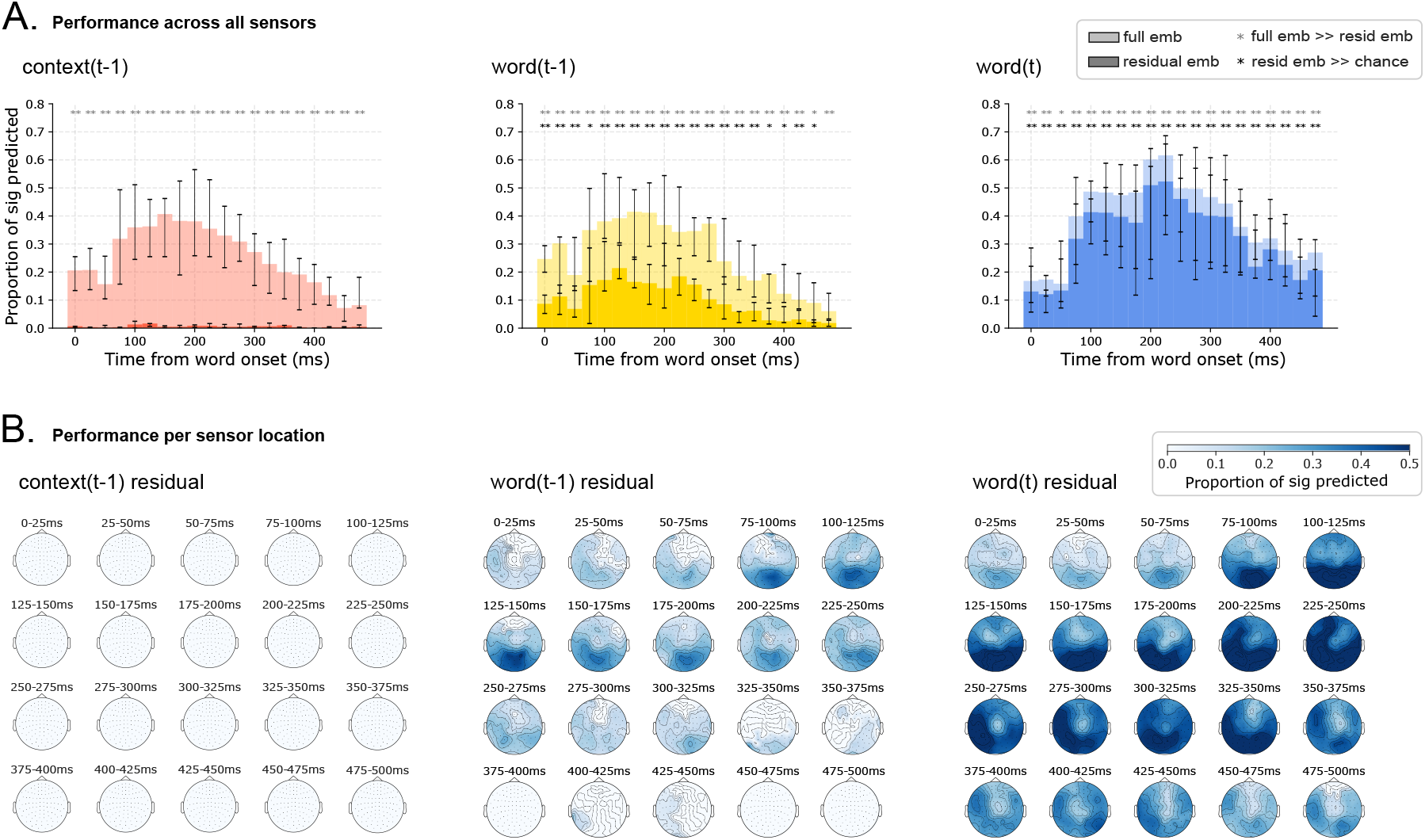
MEG prediction results at different spatial granularity. All subplots present the median across participants and errorbars signify the medians’ 95% confidence intervals. **(A)** Proportion of sensors for each timepoint significantly predicted by the full and residual embeddings (visualized in lighter and darker colors respectively). Removing the shared information among the full current word, the previous word and the context embeddings results in a significant decrease in performance for all embeddings and lobes. The decrease in performance for the context embedding (left column) is the most drastic, with no timewindows being significantly different from chance for the residual context embedding. **(B)** Proportions of sensor neighborhoods significantly predicted by each residual embedding. Only the significant proportions are displayed (FDR corrected, *p* < 0.05). Context-residuals do not predict any sensor-timepoint neighborhood while both the previous and the current word residuals predict a large subset of sensor-timepoints, with performance peaks in occipital and temporal lobes.

To understand the source of this salience, we investigated the relationship between the MEG recordings and the word embeddings for the currently-read and previously-read words. One approach to reveal this relationship is to train an encoding model as a function of the word embedding (Jain and Huth, 2018; Toneva and Wehbe, 2019). However, the word embedding corresponding to a word at position *t* is correlated with the surrounding word embeddings (Fig. 1C). Therefore, part of the prediction performance of the word *t* embedding may be due to processing related to previous words. To isolate processing that is exclusively related to an individual word, we constructed “residual word embeddings”, following the approach of constructing the residual context embeddings (see Materials and Methods). We observe that the residual word embeddings for the current and previous words lead to significantly worse predictions of the MEG recordings, when compared to their corresponding full embeddings (Fig. 3A, middle and right panels; timepoint-level Wilcoxon signed-rank test, *p* < 0.05, Benjamini-Hochberg FDR correction). This indicates that a significant proportion of the activity predicted by the current and previous word embeddings is due to the shared information with surrounding word embeddings. Nonetheless, we find that the residual current word embedding is still significantly predictive of brain activity everywhere the full embeddings was predictive. This indicates that properties unique to the current word are well predictive of MEG recordings at all spatial granularity. The residual previous word embedding predicts fewer time windows significantly, particularly 350-500ms post word *t* onset. This indicates that the activity in the first 350ms when a word is on the screen is predicted by properties that are unique to the previous word. Taken together, these results suggest that the properties of recent words are the elements that are predictive of MEG recordings, and that MEG recordings do not reflect the supra-word meaning beyond these recent words.

Lastly, we directly compared how well each imaging modality can be predicted by each meaning embedding (Fig. 4). Residual embeddings predict fMRI and MEG with significantly different accuracy (Fig. 4A), with fMRI being significantly better predicted than MEG by the residual context, and MEG being significantly better predicted by the residual of the previous and current words (Wilcoxon rank-sum test, *p* < 0.05, Holm-Bonferroni correction). In contrast, the full context embeddings do not show a significant difference in predicting fMRI and MEG recordings(Fig. 4B). We further observe that the residual embeddings lead to an opposite pattern of prediction in the two modalities(Fig. 4C). While the residual context predicts fMRI the best out of the three residual embeddings, it performs the worst out of the three at predicting MEG (Wilcoxon signed-rank test, *p* < 0.05, Holm-Bonferroni correction). In contrast, the full context and previous word embeddings do not show a significant difference in MEG prediction (Fig. 4D), suggesting that it is the removal of individual word information from the context embedding that leads to a significantly worse MEG prediction. These findings further suggest that fMRI and MEG reflect different aspects of language processing – while MEG recordings reflect processing related to the recent context, fMRI recordings capture the contextual meaning that is beyond the meaning of individual words.

**Figure 4:**
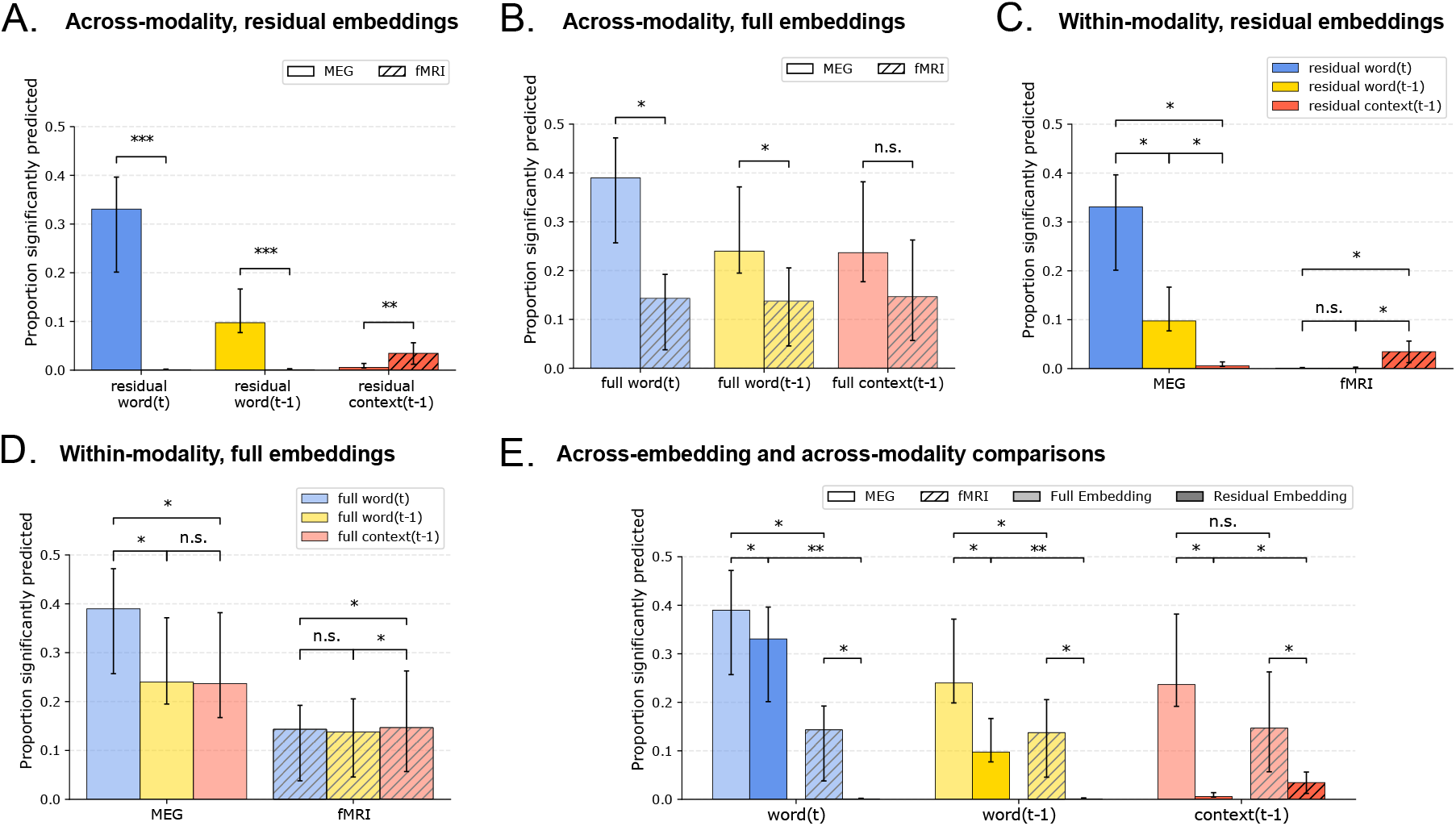
Direct comparisons of prediction performance of different meaning embeddings. Displayed are the median proportions across participants and the medians’ 95% confidence intervals. Differences between modalities are tested for significance using a Wilcoxon rank-sums test. Differences within modality are tested using a Wilcoxon signed-rank test. All p-values are adjusted for multiple comparisons with the Holm-Bonferroni procedure at *α* = 0.05. **(A)** Residual previous word, context, and current word embeddings predict fMRI and MEG with significant differences. **(B)** Full context embeddings do not predict fMRI and MEG with significant differences, while the full current word and previous word embeddings predict MEG significantly better than fMRI. **(C)** MEG and fMRI display a contrasting pattern of prediction by the residual embeddings. The current word residual best predicts MEG activity, significantly better than the previous word residual, which in turns predicts MEG significantly more than the context residual. In contrast, the context residuals significantly predict fMRI activity better than the previous and current word residuals. **(D)** Full previous word and context embeddings do not predict MEG significantly differently. **(E)** All full embeddings predict both fMRI and MEG significantly better than the corresponding residual embeddings.

## Discussion

We enabled the investigation of emergent multi-word meaning, or supra-word meaning, in the brain by devising a computational representation of it that combines representations of natural text from recent neural network algorithms with a computational control that disentangles composed-from individual-word meaning. We investigated the spatial and temporal processing signatures of supra-word meaning by evaluating its ability to predict specific locations and timepoints of recorded brain activity via fMRI and MEG respectively.

After conducting a replication analysis with a second fMRI dataset from a movie-watching paradigm and an independent set of participants, we found that our devised supra-word meaning representation consistently predicts fMRI recordings in the bilateral anterior and posterior temporal lobes (ATL and PTL). This finding supports some current hypotheses of language composition in the literature. Specifically, our results provide new evidence that the ATL processes composed meaning beyond simple concrete concepts, which supports the hypothesis that the ATL is a semantic integration hub (Visser *et al*., 2010; Pallier *et al*., 2011; Pylkkänen, 2020). Our results may also align with the hypothesis that the posterior superior temporal sulcus (pSTS, part of the PTL) is involved in building a type of supra-word meaning, by integrating information about the verb and its arguments with other syntactic information (Friederici, 2011; Frankland and Greene, 2015; Skeide and Friederici, 2016). Further, our findings pose questions for the theory that posits left PTL as primarily a site of lexical (i.e. word-level) semantics, and left IFG as a hub of integrated contextual information (Hagoort, 2020). It also poses questions for the theory that combinatorial semantics are processed in the ATL while lexical semantics are processed in more posterior regions (Hickok and Poeppel, 2007). Our finding that the PTL maintains supra-word meaning indicates that the role of the PTL extends beyond word-level semantics and suggests a common substrate for lexical and combinatorial semantics. Further, we do not find evidence for supra-word meaning in left IFG, though this does not prove that left IFG does not represent supra-word meaning – the lack of significance may be due to low statistical power. Lastly, the finding that clusters of voxels in the PTL and ATL share a common representation of composed meaning suggests that the two areas may be working together to maintain the supra-word meaning.

Strikingly, we found that, even though our devised supra-meaning representation predicted a significant proportion of fMRI voxels, it did not significantly predict any sensor-timepoints in MEG. Instead, the MEG recordings were significantly predicted by information unique to both the currently-read and previously-read words. This result was repeated when using another language model (GPT-2) to analyse the reading data and when using data from a subject listening to spoken stories. We emphasize that this null result is accompanied by two strongly significant results which, when taken together, should alleviate some concerns about the quality of the MEG data and the quality of the supra-word meaning embedding. These two strongly significant results are that: 1) the MEG data is indeed strongly predicted by representations of both individual words and context (Fig. 3), and 2) the supra-word meaning embedding predicts significant proportions of the fMRI recordings (Fig. 2). There is a null result *only* when the supraword meaning embeddings are used to predict the MEG data. Given these results, we are confident that removing the individual word information from the adjacent context eliminates much of the information that is useful to predict the MEG recordings. These findings suggest a difference in the underlying brain processes that fMRI and MEG capture. Indeed, while it is widely known that fMRI and MEG recordings result from different physiological signals, whether they capture the same underlying brain processes is still debated (Hall *et al*., 2014). Our results suggest that fMRI recordings are sensitive to supra-word meaning, while MEG recordings reflect instantaneous processes related to both the current word being read and the previously-read word. A likely candidate for the instantaneous process reflected in MEG is the process of integrating the current word with the previous context. The sensitivity to the previously-read word has many possible explanations. One possible explanation is that a word might take longer to process and integrate into the composed meaning than the duration it is on the screen. Another possible explanation is that a word may constrain the processing of the word that follows it, highlighting its relevant properties and aiding with composition. The hypothesis that MEG recordings reflect the process of composition aligns well with a vast number of previous findings characterizing transient responses evoked by a stimulus that is difficult to integrate with the preceding context (Kutas and Federmeier, 2011; Kuperberg *et al*., 2003; Kuperberg, 2007; Rabovsky *et al*., 2018) and results showing that MEG recordings are better fit by a model constrained by the meaning of the immediately preceding words (Lyu *et al*., 2019). Indeed, our results are not in disagreement with this literature – they do not show that MEG activity does not reveal word integration processes that depend on previous context. Instead, our results suggest that the representation of that previous context is not visible in MEG.

The observed difference in predicting fMRI and MEG recordings raises the hypothesis that the process of maintenance of the composed meaning does not rely on neural mechanisms that are thought to generate the MEG signal (such as synchronized current flow in pyramidal cell dendrites (Hall *et al*., 2014)) but on some other mechanisms that are not visible in the MEG signal or might be indistinguishable from noise (e.g. unsynchronized neural firing), but that have enough metabolic demands to generate a BOLD response. One explanation may be that because the MEG signal is thought to be most related to synchronized firing of pyramidal cells, it best reflects the representation of language processes which are thought to be supported by pyramidal cells. For example, the N400 and other ERPs related to composition operations, are measurable in both MEG and EEG, and thus are likely to recruit pyramidal cells. Working memory processes have also been shown to involve pyramidal cell firing (Goldman-Rakic, 1996). While both the N400 and working memory are thought to reflect short-term transient processing (Luck *et al*., 1996; Courtney *et al*., 1997), maintaining the supra-word meaning may depend on longer-term memory processes, which may be supported by different cell types or mechanisms. Alternate possible explanations for the lack of predictability of MEG by supra-word meaning are that the representation of supra-word meaning may be too distributed to be captured by MEG due to its poor spatial resolution. However, we observe that the supra-word meaning predicts about 5 – 10% of all cortical fMRI voxels across participants, mostly centered in the ATL and PTL. Thus, it is unlikely that no MEG sensor-timepoint is sensitive to this signal if it is detectable in the magnetic field changes. Further, MEG is known to be sensitive to neural activity that originates in the sulci, and since we find that the voxels that are sensitive to supra-word meaning are in the sulci, this explanation is even less likely. Another possibility is that our MEG dataset presented a false negative due to particularities of the experimental paradigm, or statistical variability due to noise. The replication in one additional subject with a natural speech listening paradigm, while encouraging, will need to be followed by future work that considers a larger number of subjects with a comparable number of trials per subject (5000 for the Harry Potter MEG experiment), which are both important for evaluating the significance of the results (Chen *et al*., 2022). Such future work will require the use of MEG datasets where each subject listens to a natural connected speech for a long duration (at least an hour per subject). Our results, if replicated in other studies, will call for a more nuanced understanding of previous work that aims to study composition of sentence-level meaning using MEG as well as possibly other types of imaging modalities that rely on synchronized firing, such as EEG and ECoG. Our results suggest that observed increases in activity measured by these modalities during sentence reading (Fedorenko *et al*., 2016; Hultén *et al*., 2019) and improved fit by a model constrained by very recent context (Lyu *et al*., 2019) may be due to instantaneous integration processes rather than the maintenance of sentence-level meaning. Future work is needed to understand whether and how these imaging modalities can be used to study sentence-level meaning.

According to our definition, every sentence has some supra-word meaning. Mathematically, we know that the supra-word meaning embedding (i.e. the embedding calculated by context(t-1)-g(word(t), word(t-1),..word(t-n)) contains information because context(t-1) is a nonlinear function of word(t-1),..word(t-n), whereas g(word(t), word(t-1),..word(t-n)) is a linear function. Whether that supra-word meaning is brain-relevant is not known, and this is what we test in the current work. Our results show that the supra-word meaning embedding contains brain-relevant information, because the supra-word meaning embedding predicts significant proportions of several language regions (Fig. 2B). In addition, we show that the supra-word meaning embedding contains multiple facets of brain-relevant meaning–one facet that is predictive of the bilateral ATL and the left pSTS, and the other of the right PTL. We further show that while the full context embedding from the first hidden layer of ELMo evaluated at word *t* is strongly linearly related to the next word t+1, current word t, and the previous word t-1, the supra-word embedding for word t does not strongly depend on any individual word, as it was designed to limit contributions of individual words (Suppl. Fig. S1). The presence of the strong linear relationship between the full context representation and the closely adjacent words means that any linear prediction of brain recordings may be overwhelmingly related to these adjacent words.

Pinpointing what linguistic or psychological information relates to the significant prediction of the bilateral ATL and PTL that we observe using the supra-word embeddings is an important question for future work. However, we want to emphasize that even if we understand that information X, Y, and Z is contained in the supra-word embeddings, that *will not* answer this question. The reason is that the supra-word embedding contains X, Y, Z, and possibly other information W, that we have not measured, and any one of them may be the information that is predictive of the bilateral ATL and PTL. For example, as our work shows, the MEG recordings are predicted by the full context embedding, but not by the information in the full context embedding that is orthogonal to the individual word embeddings (i.e. supra-word embeddings). Therefore, only knowing 2 things—1) full context embeddings predict MEG, and 2) some supra-word information is contained in the full context embeddings—is not enough to reveal what information in the full context embeddings is predictive of the MEG recordings, and suggesting that it may be because of the supra-word information would be misleading. Similarly, linguistic information X,Y, and Z may be contained in the supra-word embedding, but may not be necessary for predicting the bilateral ATL and PTL. This is in fact one of the central points of this work, and we believe it is critical to make this point as it is becoming increasingly popular to relate brain recordings to complex neural network-derived embeddings that contain multiple sources of information (Toneva *et al*., 2021). In this work, we show one way forward from this by utilizing computational controls. If linguistic information X,Y, and Z are shown to be contained in the supra-word embedding, one could regress all combinations of the corresponding linguistic labels from the supra-word embedding and observe how the prediction of fMRI recordings in the bilateral ATL and PTL changes as a result. These analyses are not simple and they would require the stimulus dataset to be annotated with various linguistic and psychological labels, some of which may require expert linguists who might even disagree amongst themselves. We believe this is an important undertaking and certainly a next step in this research direction of benefiting from recent progress in large neural network models while also granting us more control over the scientific inferences we can make.

Our analysis depends on the degree to which the computational neural network we have chosen is able to represent composed meaning. Based on ELMo’s competitive performance on downstream tasks (Peters *et al*., 2018) and ability to capture complex linguistic structure (Tenney *et al*., 2019), we believe that ELMo is able to extract some aspects of composed meaning. In addition, using a different language model (GPT-2) to extract the supra-word meaning also significantly predicts the bilateral ATL and PTL. The supra-word meaning obtained from GPT-2 additionally predicts the bilateral Angular Gyrus (AG) and Posterior Cingulate (PC). This added predictive power may be due to several factors that we cannot control without training a model from scratch: GPT-2 has larger embeddings (768 vs 512 for the forward LSTM in ELMo), has more hidden layers (12 vs 2 in ELMo), was pretrained on more data (40GB of text data vs 11GB of text data for ELMo), and has an all-together different architecture (Transformer-based vs recurrence-based for ELMo). Overall, evaluating the ability of different architectures to encode supra-word meaning is an interesting question, but it also needs to be approached with care due to these many differences. The degree to which the composed meaning in NLP models reflects the one in the brain is an important question that we have only begun to study and needs further investigation. Secondly, our residual approach accounts only for the linear dependence between individual word embeddings and context embeddings. By construction, the internal state of the LSTM in ELMo contains non-linear dependencies on the input word vector and the previous LSTM state. It is possible however that some dimensions of the internal state of the ELMo LSTM corresponds to non-linear operations on the dimensions of the input vector alone, without a contribution from the previous internal state of the LSTM (see Materials and Methods for the LSTM equations). This non-linear transformation of the input word might not be removed by our residual procedure, and whether it aligns with processing of individual words in the brain is a question for future research.

The surprising finding that supra-word meaning is difficult to capture using MEG has implications for future neuroimaging research and applications where natural language is decoded from the brain. While high temporal imaging resolution is key to reaching a mechanistic level of understanding of language processing, our findings suggest that a modality other than MEG may be necessary to detect long-range contextual information. Further, the fact that an aspect of meaning can be predictive in one imaging modality and invisible in the other calls for caution while interpreting findings about the brain from one modality alone, as some parts of the puzzle are systematically hidden. Our results also suggest that the imaging modality may impact the ability to decode the contextualized meaning of words, which is central to brain-computer interfaces (BCI) that aim to decode attempted speech. Recent success in decoding speech from ECoG recordings (Makin *et al*., 2020) is promising, but needs to be evaluated carefully with more diverse and naturalistic stimuli. Using BCI to decode speech in real life is complicated by the inherent uncertainty in decoding each word and the fact that the space of all possible utterances is not constrained. It is yet to be determined if word-level information conveyed by electrophysiology will be enough to decode a person’s intent, or if the lack of supra-word meaning should be compensated in other ways.

## Materials and Methods

### fMRI data and preprocessing: reading a chapter of a book

We use fMRI data of 9 participants reading chapter 9 of *Harry Potter and the Sorcerer’s Stone* (Rowling, 2012), collected and made available online by Wehbe *et al*. (2014b). Words were presented one at a time at a rate of 0.5s each. fMRI data was acquired at a rate of 2s per image, i.e. the repetition time (TR) is 2s. The images were comprised of 3 × 3 × *3mm* voxels. The data for each participant was slice-time and motion corrected using SPM8 (Kay *et al*., 2008), then detrended and smoothed with a 3mm full-width-half-max kernel. The brain surface of each participant was reconstructed using Freesurfer (Fischl, 2012), and a grey matter mask was obtained. The Pycortex software (Gao *et al*., 2015) was used to handle and plot the data. For each participant, 25000 – 31000 cortical voxels were kept.

### fMRI data and preprocessing: watching a full-length movie

We replicated our fMRI findings in a second fMRI dataset, which is provided by the Courtois NeuroMod group (data release `cneuromod-2020`). In this dataset, 6 healthy participants view the movie Hidden Figures in English. In total, approximately 120 minutes of data were recorded per participant during 12 scans of roughly equal length. The fMRI sampling rate (TR) was 1.49 seconds. The data was prepossessed using fMRIPrep 20.1.0 (Esteban *et al*., 2018). Three participants are native French speakers and three are native English speakers. All participants are fluent in English and report regularly watching movies in English. This data is available by request at https://docs.cneuromod.ca/en/latest/ACCESS.html.

### MEG data and preprocessing

The same paradigm was recorded for 8 participants using MEG by the authors of (Wehbe *et al*., 2014a) and shared upon our request. This data was recorded at 306 sensors organized in 102 locations around the head. MEG records the change in magnetic field due to neuronal activity and the data we used was sampled at 1kHz, then preprocessed using the Signal Space Separation method (SSS) (Taulu *et al*., 2004) and its temporal extension (tSSS) (Taulu and Simola, 2006). The signal in every sensor was downsampled into 25ms non-overlapping time bins. For each of the 5176 word in the chapter, we therefore obtained a recording for 306 sensors at 20 time points after word onset (since each word was presented for 500ms).

### ELMo details

At each layer, for each word ELMo combines the internal representations of two independent LSTMs – a forward LSTM (containing information from previous words) and a backward LSTM (containing information from future words). We extracted context embeddings only from the forward LSTM in order to more closely match the participants, who have not seen the future words. For a word token *t*, the forward LSTM generates the hidden representation 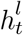 in layer *l* using the following update equations:

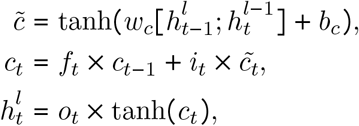

where *b_c_* and *w_c_* represent the learned bias and weight, and *f_t_, o_t_*, and *i_t_* represent the forget, output, and input gates. The states of the gates are computed according to the following equations:

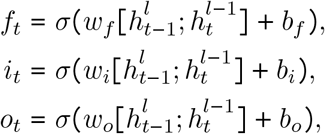

where *σ*(*x*) represents the sigmoid function and *b_x_* and *w_x_* represent the learned bias and weight of the corresponding gate. The learned parameters are trained to predict the identity of a word given a series of preceding words, in a large text corpus. We use a pretrained version of ELMo with 2 hidden LSTM layers provided by Gardner *et al*. (2018). This model was pretrained on the 1 Billion Word Benchmark (Chelba *et al*., 2014), which contains approximately 800 million tokens of news crawl data from WMT 2011.

### GPT-2 details

GPT-2 (Radford *et al*., 2019) is a transformer-based model. The pretrained GPT-2 model that we used (GPT-2 small) consists of 12 stacked transformer decoders. Unlike BERT (Devlin *et al*., 2018), GPT-2 is a causal language model, which only takes the past as input to predict the future (as opposed to both the past and the future). GPT-2 was pretrained on 8 million web pages, which were scraped using Reddit.

### Obtaining full stimulus representations

We obtain a full ELMo word embedding (as opposed to a residual word embedding) for word *w_n_* by passing word *w_n_* through the pretrained ELMo model and obtaining the token-level embeddings (i.e. from layer 0) for *w_n_*. If word *w_n_* contains multiple tokens, we average the corresponding token-level embeddings and use this average as the final full word embedding. We obtain a full ELMo context embedding for word *wn* by passing the most recent 25 words (*w*_*n*–24_, …, *w_n_*) through the pretrained ELMo model and obtaining the embeddings from the first hidden layer (i.e. from layer 1) of the forward LSTM for *w_n_*. If word w_n_ contains multiple tokens, we average the corresponding layer 1 embeddings and use this mean as the final full context embedding for word *w_n_*. We use 25 words to extract the context embedding because it has been previously shown that ELMo and other LSTMs appear to reduce the amount of information they maintain beyond 20 – 25 words in the past (Khandelwal *et al*., 2018; Toneva and Wehbe, 2019).

We follow the same technique to obtain full stimulus representations from GPT-2. We extract the word representations from the penultimate hidden layer in the network (layer 11 of 12).

### Obtaining residual stimulus representations

We obtain three types of residual embeddings for each word at position *t* in the stimulus set: 1) residual context(t-1) embedding, 2) residual word(t-1) embedding, and 3) residual word(t) embedding. We compute all three types using the same general approach of training a regularized linear regression, but with inputs *x_t_* and outputs *y_t_* that change depending on the type of residual embedding. The steps to the general approach are the following, given an input *x_t_* and output *y_t_*:

Step 1: Learn a linear function *g* that predicts each dimension of *y_t_* as a linear combination of *x_t_*. We follow the same steps outlined in the training of function *f* in the encoding model. Namely, we model *g* as a linear function, regularized by the ridge penalty. The model is trained via four-fold cross-validation and the regularization parameter is chosen via nested cross-validation.
Step 2: Obtain the residual 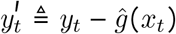, using the estimate of the *g* function learned above. This is the final residual stimulus representation.

For the residual context(t-1) embedding, the input *x_t_* is the concatenation of the full word embeddings for the 25 consecutive words *w*_*t*–24_, …,*w_t_* and the output *y_t_* is the full context(t-1) embedding. For the residual word(t-1) embeddings, the input *x_t_* is the concatenation of the full context(t-1) embedding and the full word embeddings for the 24 consecutive words *w*_*t*–24_, …, *w_t_* that exclude the full word embedding for word(t-1) and the output *y_t_* is the full word(t-1) embedding. For the residual word(t) embeddings, the input *x_t_* is the the concatenation of the full context(t-1) embedding and the full word embeddings for the 24 consecutive words *w*_*t*–24_, …, *w*_*t*–1_ and the output *y_t_* is the full word(t) embedding.

We also performed experiments with the residual context(t-1) obtained from the second hidden layer of ELMo. We did not find any significant differences in the proportion of language regions that are predicted significantly by the supra-word meaning obtained from the first hidden layer vs the supra-word meaning obtained from the second hidden layer.

### Encoding model evaluation

We evaluate the predictions of each encoding model by computing the Pearson correlation between the held-out brain recordings and the corresponding predictions in the four-fold cross-validation setting. We compute one correlation value for each of the 4 cross-validation folds and report the average value as the final encoding model performance.

### General encoding model training

For each type of embedding *e_t_*, we estimate an encoding model that takes *e_t_* as input and predicts the brain recording associated with reading the same words that were used to derive *e_t_*. We estimate a function f, such that *f*(*e_t_*) = *b*, where *b* is the brain activity recorded with either MEG or fMRI. We follow previous work (Sudre *et al*., 2012; Wehbe *et al*., 2014b,a; Nishimoto *et al*., 2011; Huth *et al*., 2016) and model f as a linear function, regularized by the ridge penalty.

The fMRI and MEG Harry Potter data is collected in 4 runs. To estimate all encoding models using this data, we perform 4-fold cross validation and the regularization parameter is chosen via nested cross-validation. Each fold holds out data corresponding to 1 run that we use to test the generalization of the estimated models. The edges of each run are removed during preprocessing (data corresponding to 20TRs from the beginning and 15TRs from the end of each run), and we further remove data corresponding to additional 5TRs between the training and test data. We follow the same procedures for the movie fMRI data, with the exception that we perform 12-fold cross validation because this dataset was recorded in 12 runs.

### fMRI Encoding Models

Ridge regularization is used to estimate the parameters of a linear model that predicts the brain activity *y^i^* in every fMRI voxel i as a linear combination of a particular NLP embedding *x*. For each output dimension (voxel), the Ridge regularization parameter is chosen independently by nested cross-validation. We use Ridge regression because of its computational efficiency and because of the results of Wehbe *et al*. (2015) showing that for fMRI data, as long as proper regularization is used and the regularization parameter is chosen by cross-validation for each voxel independently, different regularization techniques lead to similar results. Indeed, Ridge regression is indeed a common regularization technique used for building predictive fMRI (Mitchell *et al*., 2008; Nishimoto *et al*., 2011; Wehbe *et al*., 2014b; Huth *et al*., 2016).

For every voxel i, a model is fit to predict the signals 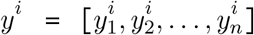, where *n* is the number of time points, as a function of the NLP embedding. The words presented to the participants are first grouped by the TR interval in which they were presented. Then, the NLP embedding of the words in every group are averaged to form a sequence of features *x* = [*x*_1_, *x*_2_, …, *x_n_*] which are aligned with the brain signals. The models are trained to predict the signal at time *t*, *y_t_*, using the concatenated vector *z_t_* formed of *[x*_*t*–1_, *x*_*t*–2_, *x*_*t*–3_, *x*_*t*–4_]. The features of the words presented in the previous volumes are included in order to account for the lag in the hemodynamic response that fMRI records. Indeed, the response measured by fMRI is an indirect consequence of brain activity that peaks about 6 seconds after stimulus onset, and the solution of expressing brain activity as a function of the features of the preceding time points is a common solution for building predictive models (Nishimoto *et al*., 2011; Wehbe *et al*., 2014b; Huth *et al*., 2016). The reason for doing this is that different voxels may exhibit different hemodynamic response functions (HRFs) so this approach allows for a data-driven estimation of the HRF instead of using the canonical HRF for all voxels.

For each given participant and each NLP embedding, we perform a cross-validation procedure to estimate how predictive that NLP embedding is of brain activity in each voxel *i*. For each fold:

- The fMRI data *Y* and feature matrix *Z* = *z*_1_, *z*_2_, …, *z_n_* are split into corresponding train and validation matrices. These matrices are individually normalized (mean of 0 and standard deviation of 1 for each voxel across time), ending with train matrices *Y^R^* and *Z^R^* and validation matrices *Y*^*V*^ and *Z^V^*.
- Using the train fold, a model *w^i^* is estimated as:

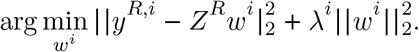 A ten-fold nested cross-validation procedure is first used to identify the best λ^*i*^ for every voxel *i* that minimizes nested cross-validation error. *w^i^* is then estimated using *λ^i^* on the entire training fold.
- The predictions for each voxel on the validation fold are obtained as *p* = *Z^V^ w^i^*.

The above steps are repeated for each of the four cross-validation folds and average correlation is obtained for each voxel *i*, NLP embedding, and participant.

### MEG encoding models

MEG data is sampled faster than the rate of word presentation, so for each word, we have 20 times points recorded at 306 sensors. Ridge regularization is similarly used to estimate the parameters of a linear model that predicts the brain activity *y^i,τ^* in every MEG sensor i at time *τ* after word onset. For each output dimension (sensor/time tuple *i,τ*), the Ridge regularization parameter is chosen independently by nested cross-validation.

For every tuple *i, τ*, a model is fit to predict the signals 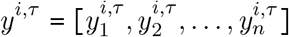, where *n* is the number of words in the story, as a function of NLP embeddings. We use as input the word vector *x* without the delays we used in fMRI because the MEG recordings capture instantaneous consequences of brain activity (change in the magnetic field). The models are trained to predict the signal at word *t*, 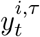, using the vector *x_t_*.

For each participant and NLP embedding, we perform a cross-validation procedure to estimate how predictive that NLP embedding is of brain activity in each sensor-timepoint *i*. For each fold:

- The MEG data *Y* and feature matrix *X* = *x*_1_, *x*_2_, … *x_n_* are split into corresponding train and validation matrices and these matrices are individually normalized (to get a mean of 0 and standard deviation of 1 for each voxel across time), ending with train matrices *Y^R^* and *X^R^* and validation matrices *Y^V^* and *Z^V^*.
- Using the train fold, a model *w*^(*i,τ*)^ is estimated as:

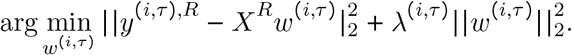 A ten-fold nested cross-validation procedure is first used to identify the best *λ*^(*i,τ*)^ for every sensor, time-point tuple (*i,τ*) that minimizes the nested cross-validation error. *w*^(*i,τ*)ℓ^ is then estimated using *λ*^(*i,τ*)^ on the entire training fold.
- The predictions for each sensor, time-point tuple (*i,τ*) on the validation fold are obtained as *p* = *X^V^ w*^(*i,τ*)^.

The above steps are repeated for each of the four cross-validation folds and an average correlation is obtained for each sensor location, time-point tuple (*s,τ*), each NLP embedding, and each participant.

### Spatial Generalization Matrices

We introduce the concept of spatial generalization matrices, which tests whether an encoding model trained to predict a particular voxel can generalize to predicting other voxels. This approach can be applied to voxels within the same participant or in other participants. The purpose of this method is to test whether two voxels relate to specific representation of the input (e.g. NLP embedding) in a similar way. If an encoding model for a particular voxel is able to significantly predict a different voxel’s activity, we conclude that the two voxels process similar information with respect to the input of the encoding model.

For each pair of voxels (*i,j*), we first follow our general approach of training an encoding model to predict voxel i as a function of a specific stimulus representation, described above, and test how well the predictions of the encoding model correlate with the activity of voxel *j*. We do this for all pairs of voxels in the PTL and ATL across all 9 participants. We finally normalize the resulting performance at predicting voxel *j* by dividing it by the performance at predicting test data from voxel *i*, that was heldout during the training process. The significance of the performance of the encoding model on voxel j is evaluated using a permutation test, described in the next subsection.

Since we are interested in semantic representation, we can formalize the tuning differences of two voxels as sensitivity to a different part of the semantic space. We can assume there exists a global, large semantic space that encodes semantic information, with the understanding that which semantic dimensions are encoded by each region, and the way in which they are encoded, can differ. We use as an approximation of the semantic space the supra-word embedding, and the way that a brain region encodes the dimensions in that space corresponds to the estimated weights of the encoding model. It is most likely that a brain region will be encoding information using a different semantic basis than our neural network derived embedding, and two brain regions might not be using the same basis. Note that the spatial generalization takes into consideration only regions that are already significantly predicted by the supra-word embedding. Since the two hypothetical regions are predicted by the encoding model using the supra-word embedding, we can assume that the supra-word embedding captures some of the dimensions spanned by their bases, up to a transformation. The difference between the bases of the two regions could manifest as some difference in the weights of the learned encoding models for the two regions. Therefore spatial generalization allows us to test how similar the semantic sensitivity of two regions are by measuring how well the estimated encoding of one voxel transfers to another voxel. Note that this is similar to methods that directly compare the encoding model weights estimated for two voxels (Çukur *et al*., 2013; Huth *et al*., 2016; Deniz *et al*., 2019), but spatial generations looks not only at how much the weights are different, but also at how much this difference affects the ability to predict held-out data. It could be considered a more stringent method, ignoring small differences in weights that don’t affect generalization performance

### Permutation tests

Significance of the degree to which a single voxel or a sensor-timepoint is predicted is evaluated based on a standard permutation test. To conduct the permutation test, we block-permute the predictions of a specific encoding model within each of the four cross-validation runs and compute the correlation between the block-permuted predictions and the corresponding true values of the voxel/sensor-timepoint. We use blocks of 5TRs in fMRI (corresponding to 20 presented words) and 20 words in MEG in order to retain some of the auto-regressive structure in the permuted brain recordings. We used a heuristic to set the block size to 5TRs–because our TR is 2 seconds, we set the block size such that it would include most of a canonical hemodynamic response, which peaks around 6 seconds and falls back to baseline over the next several seconds. We conduct 1000 permutations and calculate the number of times the resulting mean correlation across the four cross-validation folds of the permuted predictions is higher than the mean correlation from the original unpermuted predictions. The resulting p-values for all voxels/sensor-timepoints/time-windows are FDR corrected for multiple comparisons using the Benjamini-Hochberg procedure (Benjamini and Hochberg, 1995).

### Chance proportions of ROI/timewindows predicted significantly

To establish whether a significant proportion of an ROI/timewindow is predicted by a specific encoding model, we contrast the proportion of the ROI/timewindow that is significantly explained by the encoding model with a proportion of the ROI/timewindow that is significantly explained by chance. We do this for all proportions of the same ROI/timewindow across participants, using a Wilcoxon signed-rank test. We compute the proportion of an ROI/timewindow that is significantly predicted by chance using the permutation tests described above. For each permutation *k*, we compute the p-value of each voxel in this permutation according to its performance with respect to the other permutations. Next for each ROI/timewindow, we compute the proportion of this ROI/timewindow with p-values< 0.01 after FDR correction, for each permutation. The final chance proportion of an ROI/time-window for a specific encoding model and participant is the average chance proportion across permutations.

### Confidence intervals

We use an open-source package (Sheppard *et al*., 2020) to compute the 95% bias-corrected confidence intervals of the median proportions across participants. We use bias-corrected confidence intervals (Efron and Tibshirani, 1994) to account for any possible bias in the sample median due to a small sample size or skewed distribution (Miller, 1988).

### Experiments revealing shared information among NLP embeddings

For each of the 3 NLP embedding types (i.e. context(t-1) embedding, word(t-1) embedding, word(t) embedding), we train an encoding model taking as input each NLP embedding and predicting as output the word embedding for word(i), where *i* ∈ [*t* – 6, *t* + 2]. We evaluate the predictions of the encoding models using Pearson correlation, and obtain an average correlation over the four cross-validation folds.

## Data availability

Two of the three datasets analyzed during this study are included in this published article (and its supplementary information files). The remaining dataset is available by request at https://docs.cneuromod.ca/en/latest/ACCESS.html.

## Code availability

All custom scripts are included in the supplementary files of this published article, and are available without restrictions.

## Acknowledgments

The authors thank Erika Laing and Daniel Howarth for help with data collection and preprocessing, and Michael J. Tarr for helpful feedback on the manuscript. This research was supported in part by start-up funds in the Machine Learning Department at Carnegie Mellon University, the Google Faculty Research Award and the Air Force Office of Scientific Research through research grants FA95501710218 and FA95502010118.

## Author Contributions

L.W. and T.M. selected the experimental stimuli. L.W. collected the fMRI and MEG data. All authors helped conceive and design the experimental analyses and analysed the data. M.T. developed the technique to remove shared information in neural network embeddings and conducted subsequent analyses. M.T. and L.W. wrote the original draft of the manuscript. All authors contributed to the review and editing.

