## Supplementary for "Combining computational controls with natural text reveals new aspects of meaning composition"

**This PDF file includes:**

Figs. S1 to S16

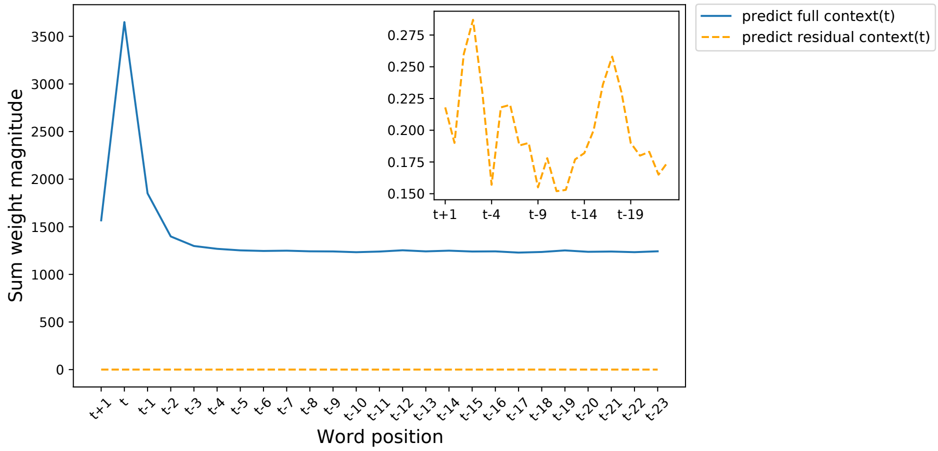


Figure S1: Importance of individual word embeddings for predicting the full and residual context embeddings. (Blue) The full context embedding from the first hidden layer of ELMo evaluated at word t is strongly linearly related to the next word t+1, current word t, and the previous word t-1. The presence of the strong linear relationship between the full context representation and the closely adjacent words means that any linear prediction of brain recordings may be overwhelmingly related to these adjacent words. In contrast, the supra-word embedding for word t (orange) does not strongly depend on any individual word, as it was designed to limit contributions of individual words.

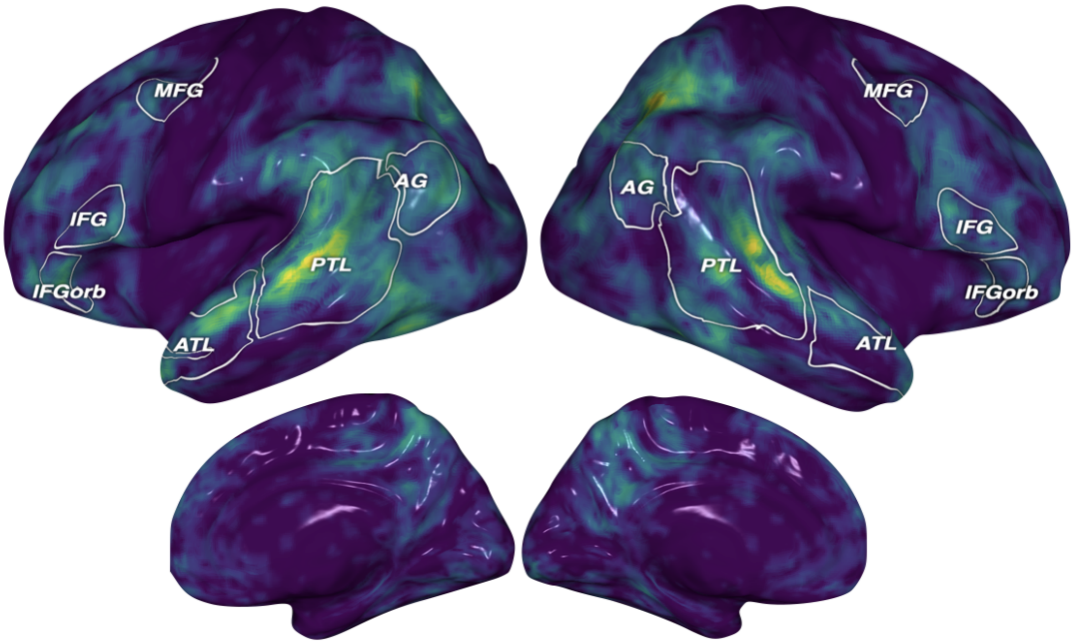


Figure S2: Harry Potter reading data: average across subjects S1-S4 shown in Fig.2 of the prediction performance using the full context embeddings. Regions that are well predicted include the bilateral posterior and anterior temporal cortices, angular gyri, inferior frontal gyri, posterior cingulate, and dorsomedial prefrontal cortex.

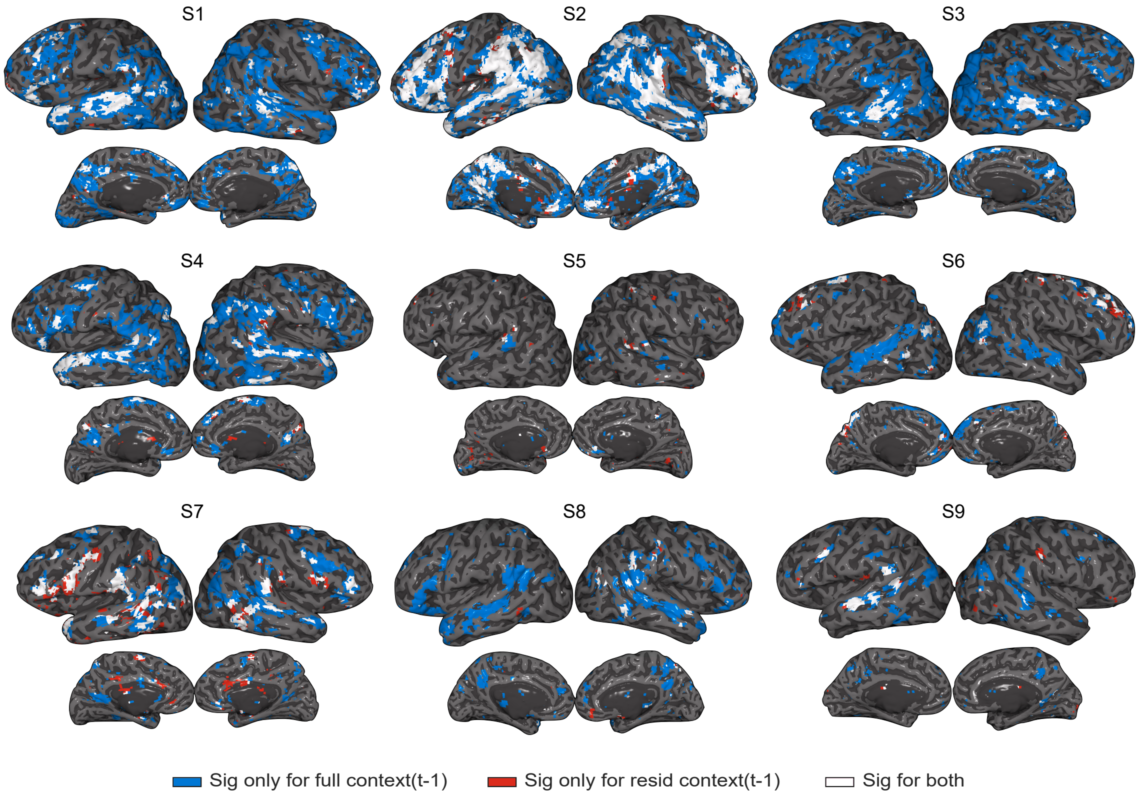


Figure S3: Harry Potter reading data: Qualitative visualization of the voxels that are significantly predicted by the full contextualized representation (in blue), the residual contextualized representation (in red), or both (in white). A voxel is determined to be significantly predicted through a permutation test and FDR correction for multiple comparisons at the 0.01 level. Large parts of the language system, spanning the temporal cortex and the inferior frontal cortex, are significantly predicted by the full context embeddings. The voxels significantly predicted by the context residual are largely a subset of those predicted by the full context embeddings.

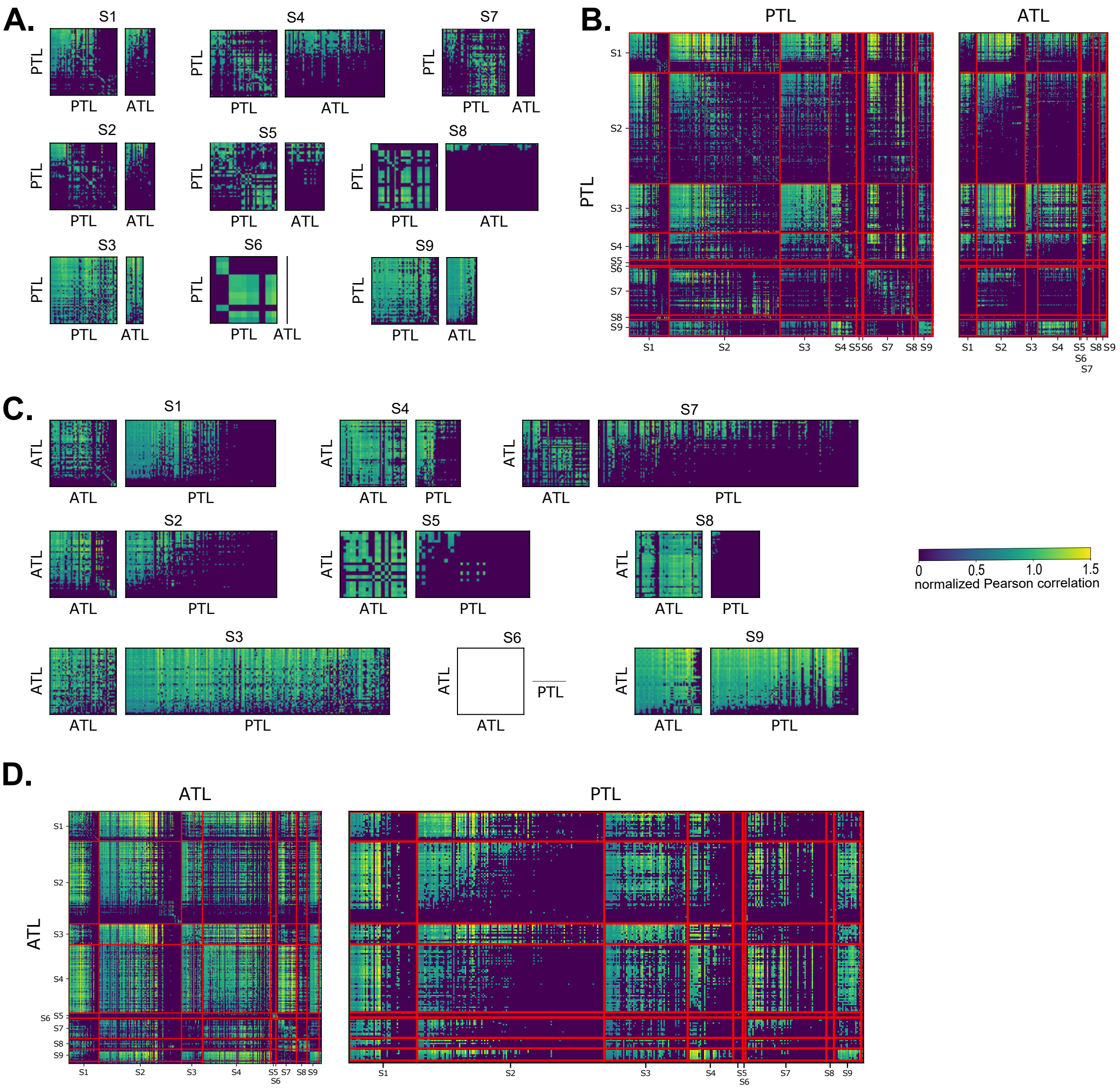


Figure S4: Harry Potter reading data: Spatial Generalization Matrices for all 9 participants. Models trained to predict PTL voxels are used to predict PTL and ATL voxels (within-participant (**A**), and across-participants (**B**)). Models trained to predict ATL voxels are used to predict ATL and PTL voxels (within-participant (**C**), and across-participants (**D**)). Note that the block diagonal matrices of the across-participants correlations (in **B**/**D**) are equivalent to the plots in **A**/**C**. Only voxels that are significantly predicted by the context residual are included in this analysis. Note that the participant S6 does not have any significantly predicted voxels in the ATL. Correlations are normalized by dividing the performance of a model trained on voxel i at predicting the target voxel j by the performance of a model trained on the target voxel j. PTL cross-voxel correlations form two clusters: voxels in a cluster can predict each other but not the other cluster's voxels. Across participants, only one of these clusters has voxels that predict ATL voxels.

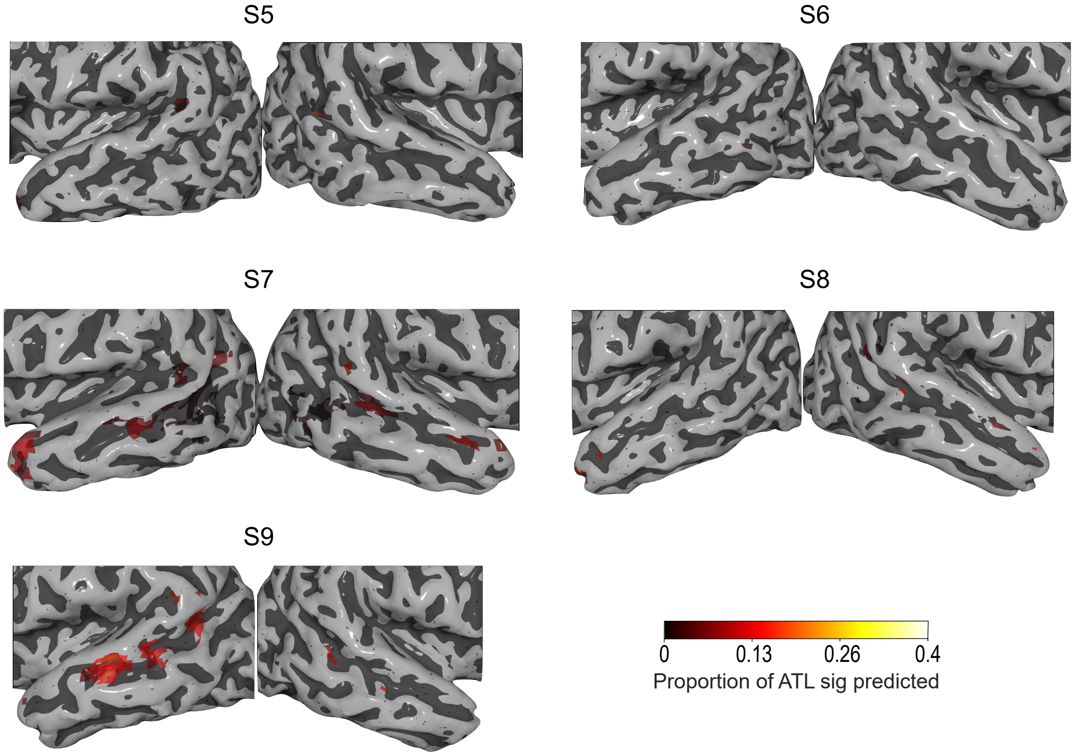


Figure S5: Harry Potter reading data: Performance of encoding models trained on ATL and PTL voxels at predicting other participants' ATL for the remaining $5$ participants. All participants who have more than a few significantly predicted voxels ($6$ out of $9$ participants) show a cluster of predictive voxels in the pSTS.

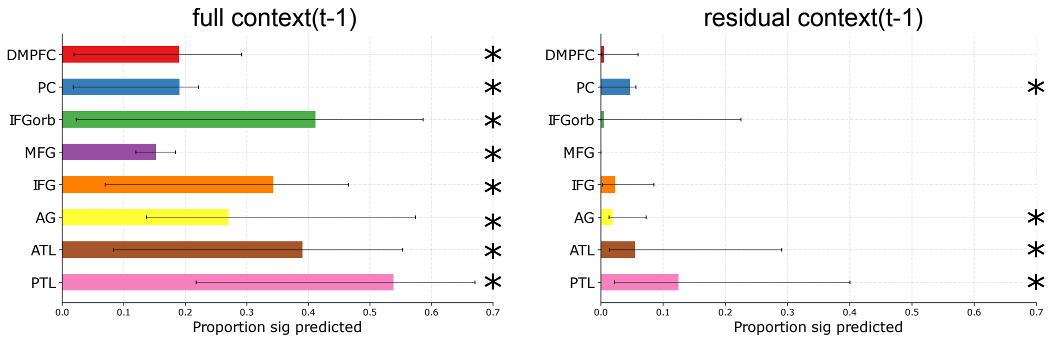


Figure S6: Harry Potter reading data: fMRI prediction results for embeddings obtained using GPT-2. Proportion of language ROI voxels significantly predicted by (Left) full context and (Right) residual context embeddings. Displayed are the median proportions across all 9 participants and the medians' 95% confidence intervals. Similarly to the full context obtained using ELMo (Fig. 2B), the full context extracted from GPT-2 predicts all ROIs (ROI-level Holm-Bonferroni correction, $p < 0.05$). The residual context predicts bilateral ATL, PTL, Angular Gyrus, and Posterior Cingulate, which replicates the results using ELMo that supra-word meaning is predictive of the bilateral ATL and PTL (Fig. 2B).

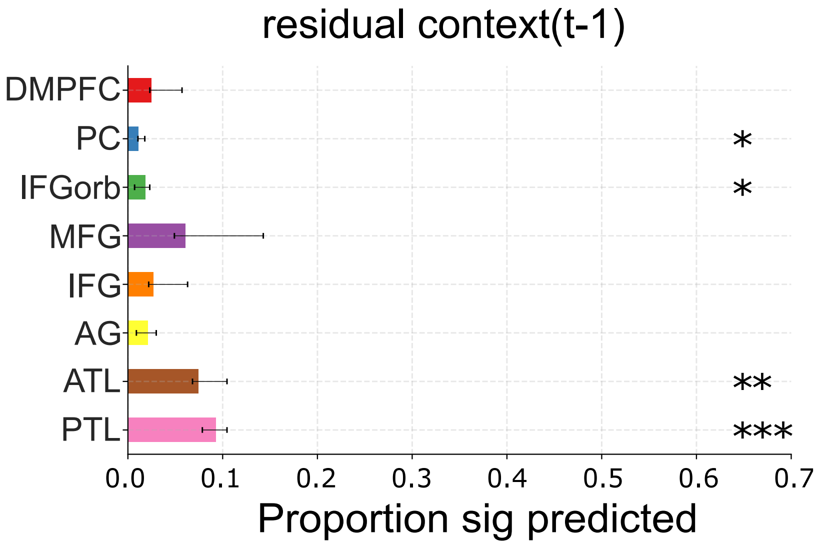


Figure S7: Hidden Figures movie data: Proportion of ROI voxels significantly predicted by residual context embeddings in the Language system ROIs (Fedorenko *et al.*, 2010) and two semantic ROIs (Binder *et al.*, 2009). Displayed are the median proportions across all participants and the medians' 95% confidence intervals. Full context predicts all ROIs (ROI-level Holm-Bonferroni correction, $p < 0.05$), while residual context predicts bilateral ATL, PTL, IFG orbitalis, and the Posterior Cingulate.

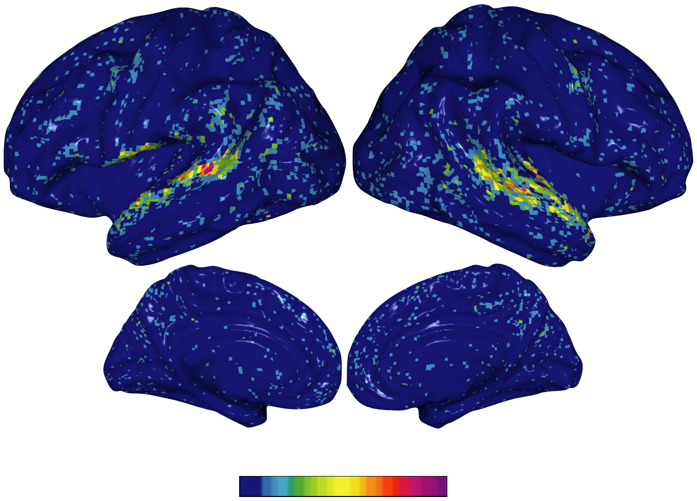


Figure S8: Hidden Figures movie data: Group-level significance map of supra-word meaning predictions in the movie fMRI data. We visualize the percentage of all 6 participants in the movie fMRI experiment for which a specific voxel is significantly predicted by ELMo's supra-word meaning (permutation test, followed by FDR correction). We observe consistent supra-word predictive performance in the bilateral ATL and PTL across participants.

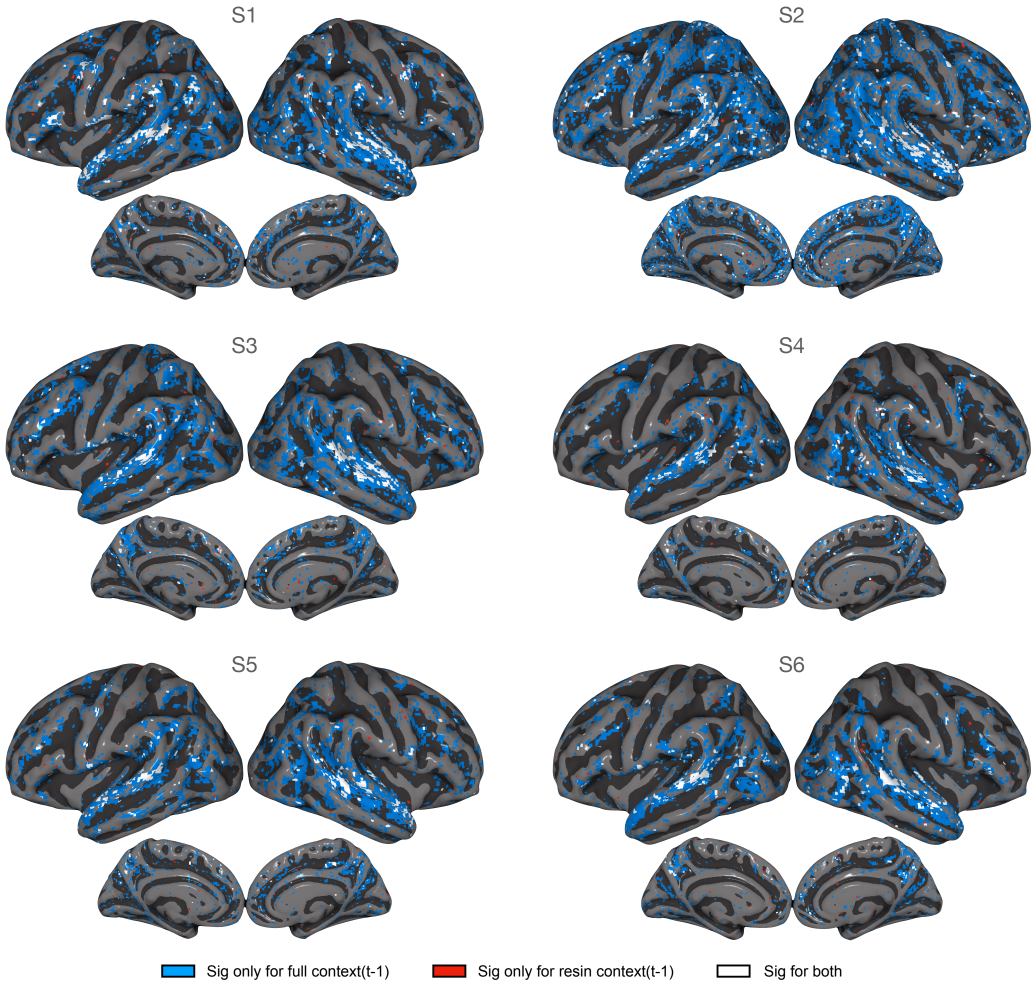


Figure S9: Hidden Figures movie data: Individual-level significance maps of prediction performance in the movie fMRI data. Voxels significantly predicted by full-context ELMo embeddings (blue), residual-context ELMo embeddings (red), or both (white), visualized in MNI space. The voxel-level significance is obtained using permutation tests and is FDR corrected. Most of the temporal cortex and IFG is predicted by full context embeddings, with residual context embeddings mostly predicting a subset of those areas.

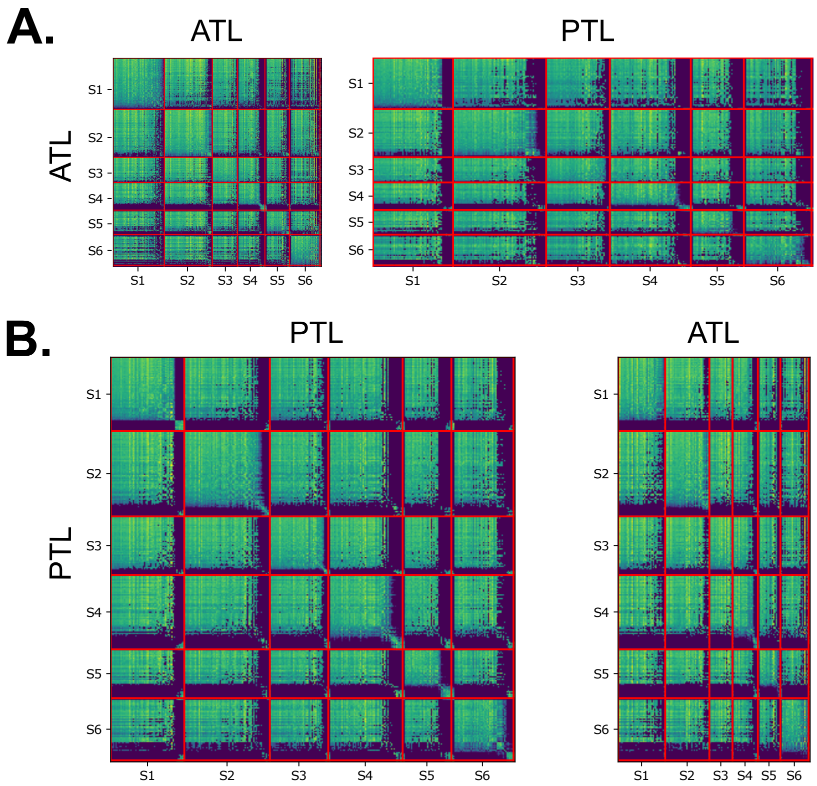


Figure S10: Hidden Figures movie data: Spatial Generalization Matrices for all 6 participants. Models trained to predict ATL voxels are used to predict PTL and ATL voxels (**A**). Models trained to predict PTL voxels are used to predict ATL and PTL voxels (**B**). Note that the block diagonal matrices of the across-participants correlations (in **A**/**B**) are equivalent to the within-participant correlations. Only voxels that are significantly predicted by the context residual are included in this analysis. Correlations are normalized by dividing the performance of a model trained on voxel i at predicting the target voxel j by the performance of a model trained on the target voxel j. PTL cross-voxel correlations form two clusters: voxels in a cluster can predict each other but not the other cluster's voxels. Across participants, only one of these clusters has voxels that predict ATL voxels.

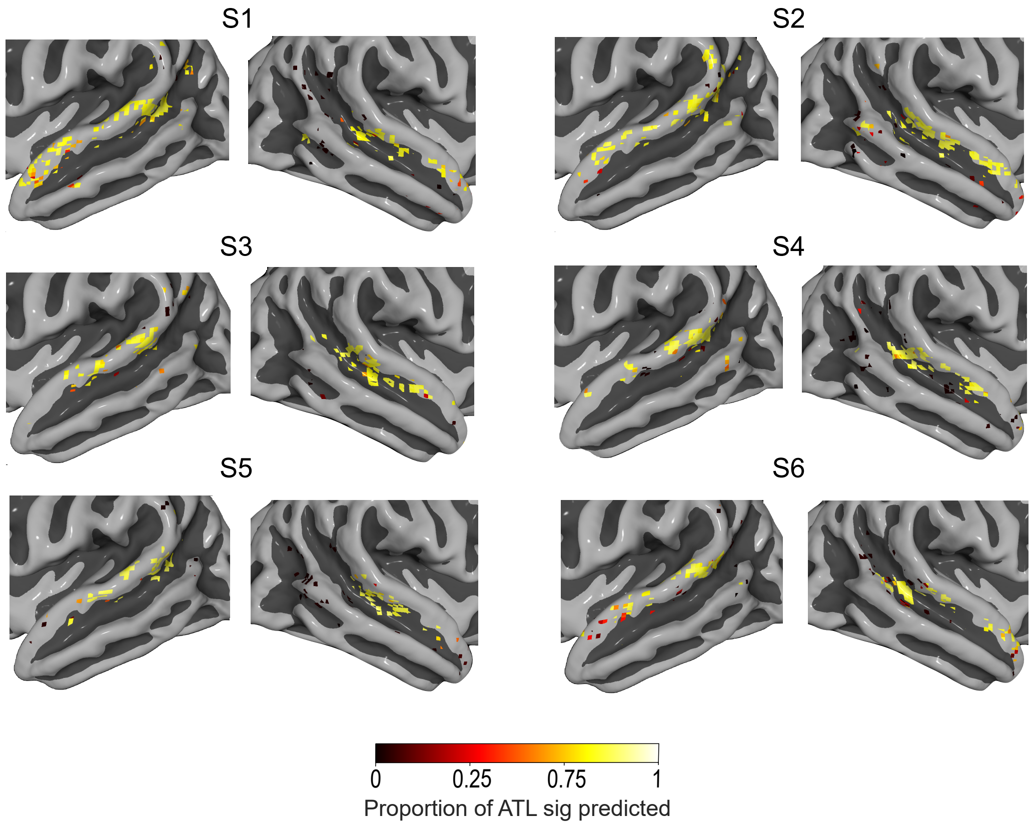


Figure S11: Hidden Figures movie data: Performance of encoding models trained on ATL and PTL voxels at predicting other participants' ATL for all 6 participants. All participants show a cluster of voxels in the left pSTS that generalize significantly to the ATL. Five out of 6 subjects show a cluster of voxels in the right pSTS that do not generalize to the ATL.

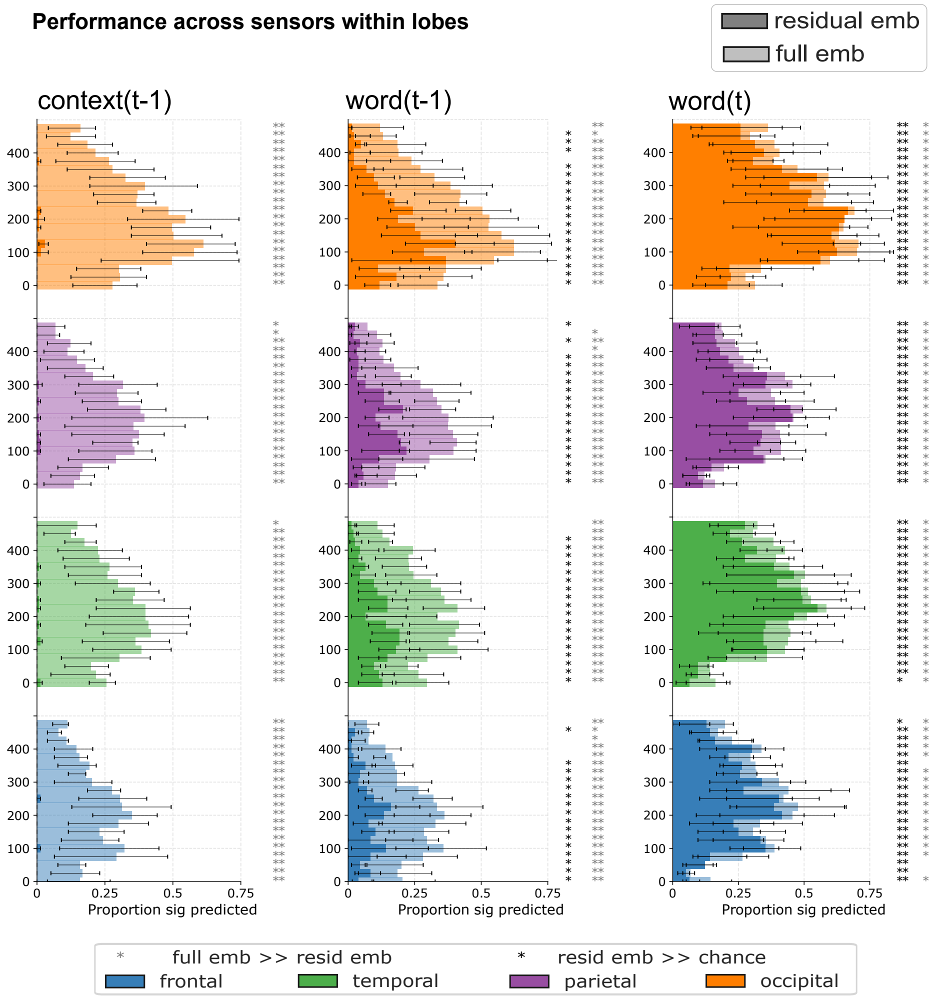


Figure S12: Proportions of significantly predicted MEG sensors for each timepoint, divided by lobe. All subplots present the median across participants and errorbars signify the medians' 95% confidence intervals. Residual embeddings performance is compared with that of full embeddings (darker and lighter colors respectively, FDR corrected, $p < 0.05$). Removing the shared information among the full current word, the previous word and the context embeddings results in a significant decrease in performance for all embeddings and lobes. The decrease in performance for the context embedding (left column) is the most drastic, with no timewindows being significant for the residual context embedding across lobes.

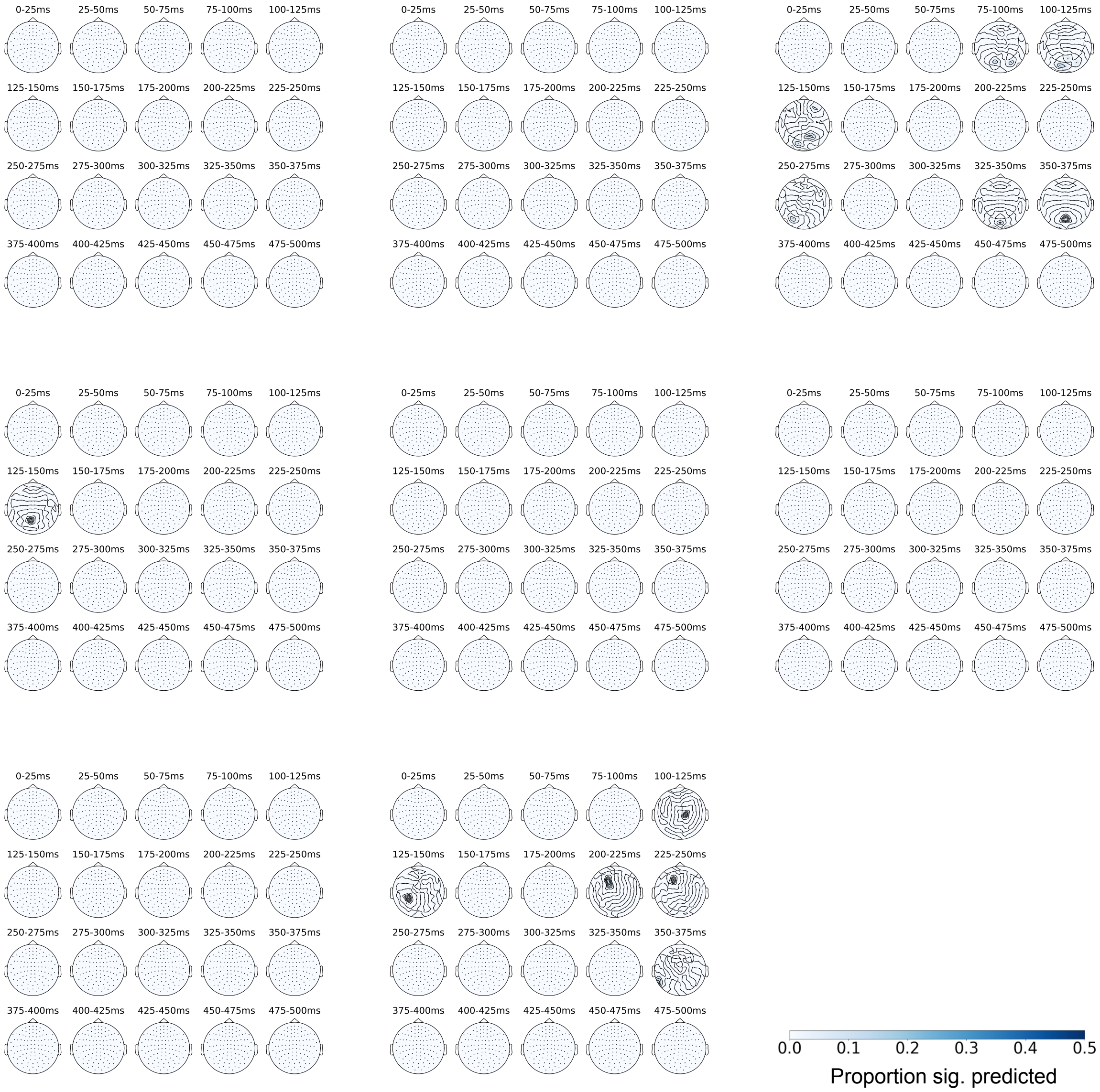


Figure S14: Individual-level proportions of MEG sensor neighborhoods significantly predicted by supra-word meaning. Only the significant proportions are displayed (FDR corrected, $p < 0.05$). Of the 6120 total MEG sensor-timepoints (306 sensors x 20 timepoints) per subject, ELMo's supra-word meaning predicts the following number significantly (at 0.05 level, permutation test followed by FDR correction): 0, 0, 1, 2, 5, 5, 20, 21. In contrast, the context predicts the following number of sensor-timepoints significantly: 3050, 4203, 3963, 1636, 3010, 5100, 4714, 4702. Removing the individual word information from the context eliminates much of the information that is useful to predict the MEG recordings.

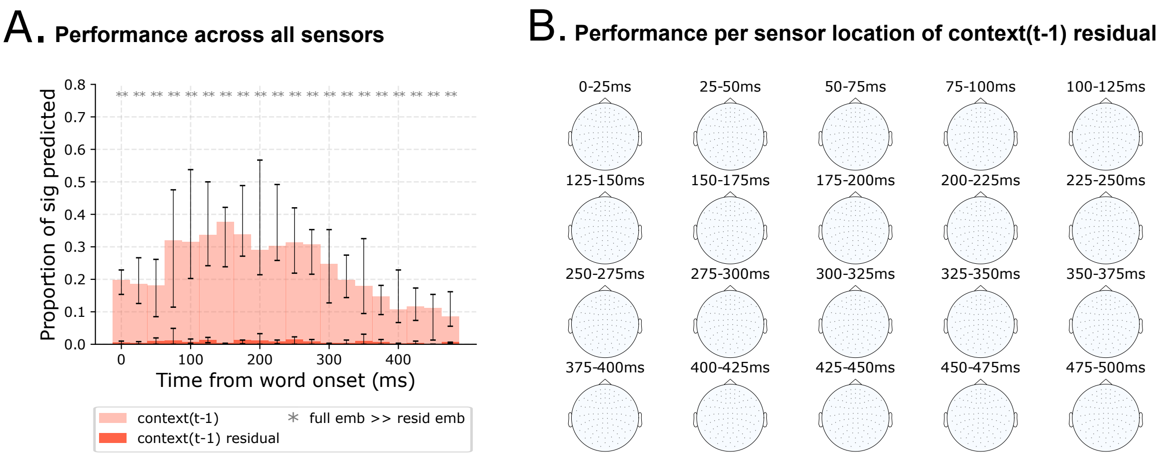


Figure S15: MEG prediction results at different spatial granularity for embeddings obtained using GPT-2. All subplots present the median across participants and errorbars signify the medians' 95% confidence intervals. **(A)** Proportion of sensors for each timepoint significantly predicted by the full and residual embeddings (visualized in lighter and darker red respectively). Removing the information related to adjacent words from the full context representation results in a significant decrease in performance for all timewindows. No timewindows are significantly different from chance for the residual context embedding. These results replicate the findings using ELMo embeddings that are presented in Fig. 3A. **(B)** Proportions of sensor neighborhoods significantly predicted by each residual embedding. Context-residuals do not predict any sensor-timepoint neighborhood significantly across participants. These results replicate the findings using ELMo embeddings that are presented in Fig. 3B.

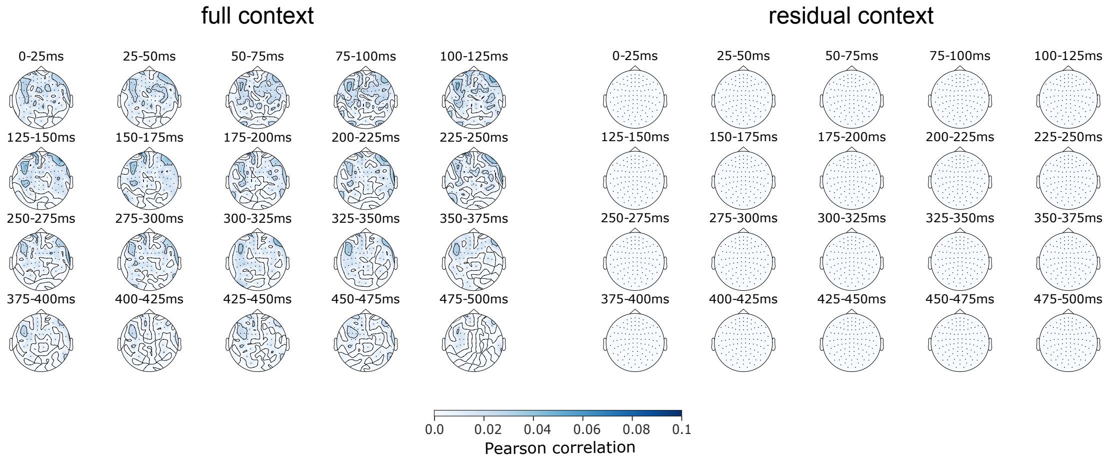


Figure S16: Prediction performance on held-out natural speech listening data when using different delays from the onset of a word. One subject listened to 70 minutes (15030 words) of stories from the moth Radio Hour while being scanned in the same Elekta machine as the one used to collect the main reading dataset (this is a replication of the Huth et al. (2016) (Huth *et al.*, 2016) dataset in MEG, and more specifically of the first of the two sessions). Onsets and offsets of words in the spoken stories were delimited by Huth et al. (2016) (Huth *et al.*, 2016). Since different words have different lengths, we build a finite impulse response model that estimates the effect of a feature at each delay, following the fMRI approach. However, we only consider one delay at a time in the model (i.e. we take the design matrix containing the embeddings of the word occurring at each time step, and delay it by the corresponding amount). This allows us to have results at a single delay, like the reading MEG results. Only the significantly predicted sensor-timepoints are displayed. [Left] Performance using the full context feature space. Many sensor/time-points, especially starting from 125ms to 350ms, are predicted with significantly higher than chance performance. The areas that are well predicted are bilateral fronto-temporal cortex, consistent with the fact that this is an auditory task that recruits the auditory cortex and the language regions, and has no visual component. We attribute the significant predictive performance of the full context in all time windows in MEG to information related to the individual words, since there is no remaining significant performance once we remove the individual word information from the full context embeddings. Specifically, the significant predictive performance in the early time windows during the presentation of word t may be related to the prediction of word t (Goldstein *et al.*, 2022), and/or to continual processing of the previous word t-1, since information about both word t-1 and word t is contained in the full context embedding. The results in Fig. 3A suggest that both types of information, independently, are predictive of these early time windows. [Right] No sensor-timepoint is predicted significantly by the residual context vector, replicating our negative results in the reading MEG data.
